# Aberrant 3′UTR splicing drives FUS-dependent mRNA condensates and prevents β-catenin from adherens junctions to promote cancer aggressiveness

**DOI:** 10.64898/2026.07.01.735936

**Authors:** Dawon Hong, Narae Kim, Yemin Jo, Jiwon Jeong, Insuk Sohn, Taeyoung Koo, Keunsoo Kang, Sunjoo Jeong

**Affiliations:** RNA Cell Biology Lab, Next-Generation RNA Editing Technology Center, Department of Bioconvergence Engineering, Dankook University Graduate School, Yongin, Republic of Korea; Arontier Inc., Gangnam-Daero 241, Seoul, Republic of Korea; Department of Pharmaceutical Sciences, College of Pharmacy, Kyung Hee University, Seoul, Republic of Korea; Department of Microbiology, Dankook University, Cheonan, Republic of Korea

**Keywords:** Alternative splicing in 3’UTR, β-catenin mRNA, mRNA localization, RNA condensates, FUS, cancer, epithelial-mesenchymal transition

## Abstract

The protein-coding sequence has long been considered the primary determinant of protein function. Alternative splicing within 3′UTRs (AS-3′UTRs) generates multiple transcript isoforms from a single gene, yet their roles in protein function and disease relevance remain largely unexplored. Through systematic transcriptome-wide identification of cancer-associated AS-3′UTRs, we uncover that AS-3′UTRs of β-catenin mRNA direct distinct subcellular localization of β-catenin mRNA isoforms. Specifically, an aberrantly spliced 3′UTR isoform promotes cytoplasmic mRNA condensate formation through FUS binding to a cancer-associated alternative exon (Exon 16A). Because β-catenin function is exquisitely dependent on its subcellular distribution between adherens junctions and the nucleus, this aberrant 3′UTR isoform reprograms β-catenin localization. By sequestering β-catenin in the cytoplasm, the aberrant 3′UTR isoform prevents its incorporation into E-cadherin-based adherens junctions, thereby inducing epithelial–mesenchymal transition (EMT)-associated transcriptional programs. Notably, the expression signature of the aberrant 3′UTR isoform robustly correlates with poor clinical outcomes in colorectal cancer patients. Together, our findings reveal that AS-3′UTRs operate as a previously unrecognized post-transcriptional regulatory mechanism through which the untranslated region of mRNA, without altering a single amino acid, reprograms protein subcellular fate to drive oncogenic phenotypes.

## INTRODUCTION

Untranslated regions (UTRs) play critical roles in post-transcriptional regulation of gene expression (1). In human cancers, widespread expression of heterogeneous 3’UTR isoforms has been reported, implicating potential roles of 3’UTR diversity in tumorigenesis (2,3). In particular, changes in 3’UTR length can profoundly influence mRNA fate, because 3’UTRs contain regulatory elements that govern mRNA stability, localization and translation (1,4). Alterations in 3’UTR lengths frequently arise from dysregulated alternative polyadenylation (5,6). In addition, alternative splicing in 3’UTR (AS-3’UTR) represents another mechanism that generates transcript diversity with variable 3’UTR length (7). AS-3’UTRs remodel UTR architecture by including or excluding exons downstream of a shared stop codon, generating multiple 3’UTR isoforms without altering protein-coding sequences (6). Consequently, AS-3′UTRs can reorganize cis-regulatory elements and RNA-binding protein (RBP) interaction sites, potentially reprogramming mRNA fate and function (8–10). Despite these observations, systematic identification and functional characterization of AS-3′UTRs remain limited, and their roles in tumorigenesis are not yet well defined (3,11).

Spatial regulation of mRNA localization and translation is essential for normal development and disease processes, particularly in controlling cell polarity and motility (12–14). Recent advances in spatial transcriptomics have enabled the mapping of subcellular mRNA localization at isoform-level resolution (15–17). Because 3′UTRs contain localization signals, distinct 3′UTR isoforms can direct differential subcellular distribution of transcripts (18). Indeed, alternative 3′UTR isoforms have been shown to regulate mRNA spatial patterns in neuronal and epithelial systems (19). RNA localization is further influenced by RBPs and RNA granules, which assemble in a context-dependent manner in response to cellular signals and stress (20,21). Among these RBPs, Fused in Sarcoma (FUS) plays a key role in RNA granule assembly and RNA trafficking (9,22,23). Under stress conditions, FUS translocates to the cytoplasm and promotes liquid-liquid phase separation (LLPS) through multivalent RNA interactions, forming RNA condensates (24–26). Although RNAs contribute to LLPS and facilitate multiphase condensate formation (27–30), whether distinct 3’UTR isoforms can modulate RNA condensate formation and spatial organization remains largely unknown.

β-catenin is a multifunctional protein that regulates both cell–cell adhesion and Wnt signaling through dynamic localization between adherens junctions and the nucleus (31–33). At the plasma membrane, β-catenin forms adherens junction complexes with E-cadherin, thereby maintaining epithelial integrity and suppressing tumor progression (34,35). Activation of Wnt signaling or mutations in *CTNNB1* (encoding b-catenin) promote cytoplasmic accumulation and nuclear translocation of β-catenin, leading to transcriptional activation of target genes involved in tumor initiation and progression (36–39). Thus, precise spatial regulation of β-catenin localization is critical for its diverse cellular functions and oncogenic potential (33,40). Although disruption of adherens junction is a hallmark of epithelial–mesenchymal transition (EMT) initiation, the upstream mechanisms regulating β-catenin incorporation into E-cadherin-based junctions remain incompletely understood. Notably, *CTNNB1* transcripts generate multiple AS-3′UTR isoforms, including cancer-associated aberrantly spliced 3′UTR variants (11,41,42). These observations strongly implicate that AS-3′UTRs regulate β-catenin localization and function through RNA-mediated spatial mechanisms.

Here, we systematically identified human genes harboring AS-3′UTRs through computational classification of intron-containing 3′UTRs. Using β-catenin mRNA as a model, we demonstrate that an aberrantly spliced 3′UTR isoform promotes FUS-dependent cytoplasmic RNA condensate formation and drives aberrant cytoplasmic localization of b-catenin without altering coding sequence. This aberrant targeting prevents β-catenin incorporation into E-cadherin-based adherens junctions and induces EMT-associated transcriptional programs. Together, our findings establish aberrant AS-3′UTRs as a post-transcriptional mechanism that reprograms protein localization through mRNA condensates to drive cancer progression.

## METHODS

### Cell culture

HCT116, HT-29, and SW480 human colorectal cancer cell lines were purchased from ATCC, and GP2-293 cells were obtained from Clontech. HCT116, HT-29, and SW480 cells were maintained in RPMI-1640 medium (Welgene) containing 4,500 mg/L glucose, 10% heat-inactivated fetal bovine serum, 100 U/ml penicillin, and 100 μg/ml streptomycin at 37°C in a humidified incubator with 5% CO₂. GP2-293 cells were cultured in DMEM supplemented with 10% fetal bovine serum and 100 U/ml penicillin and 100 μg/ml streptomycin. Cells were routinely tested for mycoplasma contamination and periodically treated with BM-Cyclin (Roche) to eliminate mycoplasma contamination.

### Generation of stable cell lines

For β-catenin overexpression experiments, stable cell lines expressing β-catenin 3′UTR isoforms were generated. pBABE-β-catenin-3′UTR-0, pBABE-β-catenin-3′UTR-1, and pBABE-β-catenin-3′UTR-2 constructs were co-transfected with the packaging vector pMD2.G (Addgene) into GP2-293 cells for VSV-G-pseudotyped retrovirus production. Viral supernatants were collected 48 h after transfection. HT-29, SW480, and SW480 β-catenin knockout cells were infected with retroviral supernatants in the presence of 8 μg/ml polybrene. At 24 h post-infection, puromycin was added at a final concentration of 0.5 μg/ml for selection. Stable cell lines were maintained in medium containing 0.5 μg/ml puromycin, and stable expression was confirmed by immunoblotting.

### CRISPR/Cas9-mediated β-catenin knockout

β-catenin knockout SW480 cells were generated using a CRISPR/Cas9-mediated genome editing approach. The guide RNA (gRNA) sequence targeting human CTNNB1 was 5′-TCCCACTAATGTCCAGCGTTTGG-3′. The gRNA sequence was cloned into a CRISPR expression vector. SW480 cells were co-transfected with CRISPR and Cas9 plasmids. After transfection, cells were diluted into 96-well plates to isolate single-cell b-catenin knockout clones. were isolated by limiting dilution in 96-well plates. β-catenin knockout efficiency was confirmed by immunoblotting and Sanger sequencing.

### siRNA-mediated knockdown

Cells were transfected with siRNAs (40 nM) using Lipofectamine RNAiMAX (Invitrogen) according to the manufacturer’s instructions. Cells were harvested 36 h after transfection, and knockdown efficiency was verified by immunoblotting. FUS/TLS siRNA and control siRNA-A were purchased from Santa Cruz Biotechnology. Detailed information is provided in the Supplementary Table 5.

### Retrovirus production

For retrovirus production, GP2-293 cells were transfected with pMD2.G and pBABE-puro vectors encoding empty vector, β-catenin-3′UTR-0, β-catenin-3′UTR-1, β-catenin-3′UTR-2, or β-catenin-3′UTR-3 constructs. The culture medium was replaced with fresh medium 6 h after transfection. Viral supernatants were collected 48 h after transfection and precipitated overnight at 4°C using PEG solution (4% PEG-8000 and 500 mM NaCl). Samples were centrifuged at 4,000 rpm for 30 min at 4°C, and viral pellets were resuspended in PBS.

### Stress conditions

To induce cellular stress, HCT116 and SW480 cells were treated with sodium arsenite at a final concentration of 1 mM for 30 min or with 0.4M sorbitol for 1 h.

### Immunoblotting

Cells were lysed in RIPA buffer containing 25 mM Tris-HCl (pH 7.6), 150 mM NaCl, 1% NP-40, 0.5% sodium deoxycholate, 0.1% SDS and protease inhibitor cocktail. Cell lysates were clarified by centrifugation at 13,000 rpm for 15 min at 4°C. Protein concentrations were measured using the Bradford assay. Protein samples were mixed with Laemmli sample buffer, boiled for 5 min, separated by SDS-PAGE, and transferred to polyvinylidene difluoride (PVDF) membranes (Millipore). Membranes were blocked with 5% non-fat milk in TBST and incubated with primary antibodies overnight at 4°C followed by HRP-conjugated secondary antibodies for 1 h at room temperature. Immunoblot signals were detected using chemiluminescence reagents. Immunoblot images are representative of at least two independent experiments.

### RNA isolation and quantitative real-time PCR

Total RNA was extracted from the cells by using the Quick-RNA MiniPrep Plus kit (Zymo Research) according to the manufacturer’s instructions. cDNA was synthesized from 2 mg total RNA using reverse transcriptase and manufacturer’s instructions and random hexamer primers (Thermo Fisher). Quantitative real-time PCR was performed PowerUp ^TM^ SYBR Green Master Mix (Thermo Fisher) on a StepOne Plus Real-time PCR system (Applied Biosystems). Relative expression levels were normalized to GAPDH expression using the ΔΔCt method.

### Immunofluorescence staining and confocal microscopy

Cells were cultured in four-well or eight-well chamber slides (LAB-TAK). Cells were washed with phosphate-buffered saline (PBS), fixed with 3.7% formaldehyde (Biosesang) for 10 min at 4°C, and permeabilized with 0.1% Triton X-100 in PBS or methanol depending on the experimental conditions. Samples were blocked with 5% goat serum in PBS for 1 h at room temperature and incubated with primary antibodies overnight at 4°C. After washing three times with PBST (PBS containing 0.1% Tween-20), samples were incubated with Alexa Fluor 488-or 547-conjugated secondary antibodies (Thermo Fisher) for 1 h at room temperature. Slides were mounted using VECTASHIELD® Antifade Mounting Medium containing DAPI. Images were acquired using an Olympus Fluoview FV300 confocal microscope equipped with a 60× oil-immersion objective lens. Primary antibodies used in this study are listed in the Supplementary Table 5.

### RNAscope in situ hybridization

RNAscope in situ hybridization was performed using the RNAscope assay (Advanced Cell Diagnostics) according to the manufacturer’s instructions. Briefly, cells were fixed with 4% paraformaldehyde and treated with protease to permeabilize the samples. β-catenin mRNA-specific probes were hybridized, followed by signal amplification using a branched DNA system. Fluorescent signals acquired using fluorescence microscope, and signal intensity or puncta per cell were quantified using ImageJ software.

### Combined RNA-FISH and IF

We performed as described previously (de Planell-Saguer et al., 2010) with minor modifications for immunostaining combined with in situ RNA visualization. RNA-FISH was performed as described above. Blocking buffer, PBS, and RNase inhibitors are provided in a kit. After the last wash in 1x PBS for 5 minutes at room temperature, the immunofluorescence protocol was followed as described above. Still, all incubations were performed in a blocking buffer with the addition of an RNase inhibitor, and all the wash steps were performed in RNase-free 1x PBS with an RNase inhibitor. Images were acquired on the Fluoview FV300 Olympus confocal microscopy with a 100x oil immersion objective lens. Image visualization and analysis were performed with ImageJ software.

### RNA Immunoprecipitation

RNA immunoprecipitation (RIP) was performed as described previously (44) with minor modifications. For UV-RIP, cells were crosslinked with ultraviolet irradiation at 0.2 J/cm². For formaldehyde-RIP, cells were fixed with 0.1% formaldehyde for 10 min at room temperature and quenched with 0.125 M glycine for 10 min. Cells were washed with ice-cold PBS and lysed in RIP lysis buffer containing 10 mM Tris-HCl (pH 8.0), 100 mM NaCl, 1 mM EDTA, 0.1% sodium deoxycholate, 0.5% SDS, 1 mM PMSF, protease inhibitor cocktail, and 100 U/ml RNaseOUT inhibitor. Lysates were treated with DNase I (400 U/ml) for 30 min at 37°C and fragmented by sonication. After centrifugation at 13,000 rpm for 15 min at 4°C, aliquots of the supernatants were collected as input controls. Remaining lysates were diluted five-fold with IP dilution buffer containing 20 mM Tris-HCl (pH 8.0), 150 mM NaCl, 1 mM EDTA, and 1% Triton X-100, followed by incubation overnight at 4°C with 2 μg mouse IgG (Thermo Fisher) or anti-FUS antibody (Abcam). Protein A/G agarose beads were added and incubated for 2 h at 4°C. Beads were sequentially washed with low-salt wash buffer, high-salt wash buffer, lithium wash buffer, and TE buffer. Bound complexes were eluted in elution buffer (1% SDS and 100 mM sodium bicarbonate) for 30 min at 65°C. RNA–protein complexes were reverse-crosslinked in proteinase K solution containing 800 μg/ml proteinase K for 1 h at 65°C. RNA was extracted using TRIzol/chloroform and purified using the Quick-RNA Miniprep Kit (Zymo Research). Purified RNA was analyzed by quantitative real-time PCR using SYBR Green Master Mix (Promega).

### Immunoprecipitation

Cells cultured in 10-cm dishes were washed twice with ice-cold PBS and harvested by centrifugation at 1,000 rpm for 5 min. Cell pellets were lysed in lysis buffer containing 20 mM Tris-HCl (pH 8.0), 150 mM NaCl, 1% NP-40, 2 mM EDTA, 250 μM PMSF, protease inhibitor cocktail (Sigma), 1 mM Na3VO4, and phosphatase inhibitor cocktail (Sigma) for 10 min on ice. Lysates were clarified by centrifugation at 13,000 rpm for 15 min at 4°C. Equal amounts of lysates (1 mg) were incubated overnight at 4°C with anti-FLAG antibody (Sigma-Aldrich, M2). Immune complexes were collected using Protein A/G beads, washed three times with lysis buffer, and boiled in 2× Laemmli sample buffer for 10 min. Eluted proteins were analyzed by immunoblotting.

### Subcellular fractionation

Subcellular fractionation was performed as previously described (45) with minor modifications. Briefly, cells (2 × 10^7^ cells/ml) were resuspended in ice-cold cytosolic buffer containing 10 mM HEPES-KOH (pH 7.5), 10 mM KCl, 1.5 mM MgCl₂, 1 mM DTT, 0.5% NP-40 and protease and phosphatase inhibitor cocktail (Sigma), followed by incubation on ice for 15 min. After centrifugation at 430 × g for 10 min at 4°C, the supernatant was collected as the cytoplasmic fraction. The remaining pellet was resuspended in nuclear buffer containing 20 mM Tris-HCl (pH 7.5), 400 mM NaCl, 2 mM EDTA, 1% Triton X-100, 0.5% sodium deoxycholate, and protease and phosphatase inhibitor cocktail, incubated on ice for 10 min, and centrifuged at 950 × g for 10 min at 4°C. The supernatant was collected as the nuclear fraction.

### Luciferase assays

Cells were seeded in 24-well plates and grown to 80–90% confluence prior to transfection. Cells were transfected in triplicate using Lipofectamine 3000 (Invitrogen) with luciferase reporter plasmids, expression constructs, and the pRL-TK Renilla luciferase control plasmid. At 24 h after transfection, firefly and Renilla luciferase activities were measured using the Dual-Glo Luciferase Assay System (Promega, E2920), and luminescence was detected using a GloMax 20/20 luminometer (Promega). Firefly luciferase activity was normalized to Renilla luciferase activity for each sample.

### Gelatin zymography

Conditioned media collected from stable cell lines were mixed with non-reducing sample buffer containing 2% SDS and 2% glycerol without β-mercaptoethanol. Samples were separated on 7.5% SDS-PAGE gels containing 1 mg/ml gelatin (Sigma-Aldrich). Gels were washed three times in renaturation buffer containing 50 mM Tris-HCl (pH 7.6), 150 mM NaCl, 10 mM CaCl₂, and 5 mM ZnCl₂, followed by incubation overnight at 37°C. Gels were stained with Coomassie Brilliant Blue and destained prior to imaging.

### Transwell invasion assay

Transwell invasion assays were performed as previously described (46) with minor modifications. Cells were resuspended in serum-free medium and seeded into the Matrigel-coated Transwell inserts (8.0 μm pore size; Corning). Medium containing 10% FBS was added to the lower chamber as a chemoattractant. After 48 h, invaded cells were fixed with 4% paraformaldehyde, stained with hematoxylin and eosin (H&E; Merck), and quantified under a light microscope.

### Survival analysis

Differentially expressed genes (DEGs) were grouped into four clusters using K-means clustering (K = 4). Gene clusters were analyzed for patient survival using GEPIA2 (47). Survival analyses were performed using The Cancer Genome Atlas colon adenocarcinoma (TCGA-COAD) cohort, including overall survival (OS) and disease-free survival (DFS). Patients were stratified into high- and low-expression groups using predefined percentile cutoffs (high: top 77%; low: bottom 16%) based on the expression level of each gene cluster. Kaplan–Meier survival curves were generated, and statistical significance was assessed using the log-rank (Mantel–Cox) test. Hazard ratios were calculated using the Cox proportional hazards model (48).

### Quantification and statistical analysis

Data are presented as mean ± SEM. Statistical significance was determined using unpaired Student’s t-test or one-way ANOVA followed by post hoc analysis, as indicated in the figure legends. A p-value < 0.05 was considered statistically significant. Unless otherwise indicated, all experiments were performed using independent biological replicates derived from separate cell cultures. Statistical analyses were performed using GraphPad Prism version 7.01.

### Differential expression analysis

Differential expression analysis was performed using DESeq2 based on raw read counts. Differentially expressed genes (DEGs) were identified from comparisons between FLAG-β-catenin-3′UTR construct-expressing SW480 cells or FLAG-β-catenin-3′UTR construct-expressing SW480 β-catenin knockout cells and corresponding control cells. Genes with an adjusted p-value < 0.05 and |log2 fold change| > 1 were defined as differentially expressed genes. Complete DEG lists and RNA-seq analysis results are provided in the Supplementary Tables.

### Gene Ontology (GO) enrichment analysis

Gene Ontology (GO) enrichment analysis was performed using Metascape (53). Gene sets selected from differential expression analysis or other predefined criteria were used for enrichment analysis.

## RESULTS

### Cancer-associated alternative splicing within 3′UTRs is widespread across the human transcriptome

To systematically identify AS-3’UTR events across the human transcriptome, we searched for transcripts containing internal introns within annotated 3’UTRs and found 2,703 genes (11.8%) harboring such splicing events (Fig. 1A, Supplementary Fig. S1A, Supplementary Table 1A). These genes were categorized into three classes based on the C-terminus of proteins and the 3’ end of mRNAs: Class I (alternative stop codon), Class II (alternative polyadenylation), and Class III (alternative splicing) (Fig. 1B). Among these, we focused on Class III comprising 320 transcript pairs from 121 genes that share identical coding sequences but differ in their 3’UTR isoforms (Fig. 1B and Supplementary Table 1B).

**Fig. 1.**
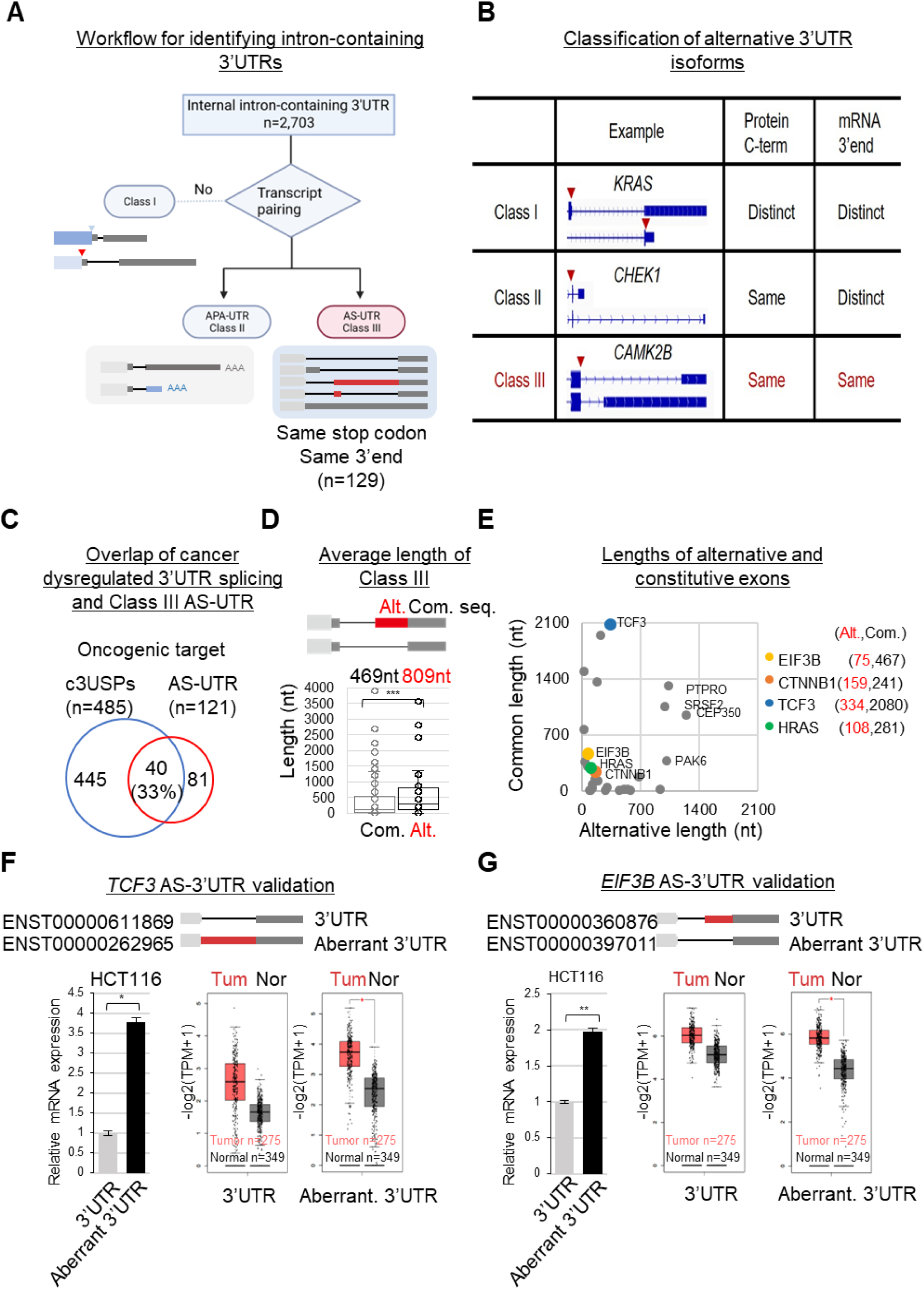
Genome-wide identification of cancer-associated alternative splicing within 3′UTRs. **(A)** Schematic overview of the pipeline used to identify intron-containing 3′UTRs in the human genome. Three classes of alternative 3′UTRs were defined: Class I, alternative stop codon (ASC); Class II, alternative polyadenylation (APA); and Class III, alternative splicing within 3’UTRs (AS-3′UTR). **(B)** Classification criteria based on the position of the translation termination codon or the 3′UTR end. Representative transcript models for each class are shown. **(C)** Venn diagram showing the overlap between cancer-dysregulated 3′UTR splicing events (c3USPs; cancer-associated 3’UTR splicing events), oncogenic targets, and prognosis-associated genes, highlighting enrichment of Class III (AS-3′UTR) candidates among cancer-relevant genes. **(D)** Distribution of alternative 3’UTR sequence lengths in Class III 3′UTRs grouped by splicing pattern. **(E)** Scatter plot comparing lengths of alternative versus shared 3′UTR regions for individual Class III target genes. **(F and G)** Expression of TCF3 **(F)** and EIF3B **(G)** AS-3′UTR isoforms in HCT116 cells and corresponding Kaplan-Meier analyses of overall survival in patients with colon adenocarcinoma (COAD).

To characterize alternative splicing events occurring in 3’UTRs, we first classified AS types and found that intron retention (IR; 71 genes) was the most frequent AS event, followed by alternative 3’ splice site (3’ss, 19 genes), cassette exon and mutually exclusive exons (ES, 19 genes), and 5’ splice site (5’ss, 5 genes), many of them generating more than two distinct 3’UTR isoforms (Supplementary Fig. S1B-S1D). Our approach successfully re-identified known AS-3’UTR events, such as in Calcium/Calmodulin-dependent protein kinase II beta (*CAMK2B*), thereby validating our annotation pipeline (54,55) (Fig. 1B and Supplementary Fig. S1C). Consistent with this, approximately 33% of Class III AS-3’UTR genes overlapped with the previously reported genes with dysregulated 3’UTR splicing in human cancers (11) (Fig. 1C and Supplementary Table 1c). Notably, alternative isoforms were significantly longer (average 809 nt) than common isoforms (average 469 nt) in cancer-associated transcripts, suggesting regulatory potential associated with alternative exons (Fig. 1D and 1E).

Gene ontology analysis of Class III genes revealed enrichment in pathways related to transport, organelle organization and Wnt/β-catenin signaling, implicating roles of AS-3’UTRs in tumorigenesis (Supplementary Fig. S1E). Consistent with this enrichment, WNT signaling-related genes such as *TCF3, EIF3B*, *HRAS* and *CTNNB1* were highlighted by colored markers among AS-3’UTR containing cancer-associated transcripts (Fig. 1E). To investigate the clinical relevance of these AS-3’UTRs in colorectal cancer (CRC), we examined the AS-3’UTR isoform expression in CRC cell (HCT116) and colorectal adenocarcinoma (COAD) patient samples. One AS-3’UTR isoform of *TCF3* and *EIF3B* was highly expressed as the prominent isoform in HCT116 and COAD, leading us to designate them as cancer-associated AS-3′UTRs (Fig. 1F, 1G and Supplementary Fig. S1F). Among these candidates, we focused on β-catenin (*CTNNB1*), a central regulator of WNT signaling frequently dysregulated in colorectal cancer, to investigate the functional consequences of aberrant 3′UTR splicing.

### An aberrant 3′UTR of β-catenin mRNA is upregulated in colorectal cancer and associates with poor clinical outcomes

Our genomic analysis suggested the expression of cancer-associated AS-3’UTR isoforms from the genes in WNT/β-catenin signaling. As a key factor in WNT signaling, β-catenin protein is a critical regulator of tumorigenesis in colorectal adenocarcinoma (COAD) (56). As a Class III gene, *CTNNB1* mRNAs harbor multiple representative AS-3’UTR isoforms, 3’UTR-1 (a short spliced 3’UTR isoform lacking the alternative exon, Canonical 3’UTR), 3’UTR-2 (the long isoform with an aberrantly spliced alternative exon, Aberrant 3’UTR), and 3’UTR-3 (an unspliced isoform with a retained intron in the 3’UTR) (Fig. 2A and Supplementary Fig. S2A). Using GEPIA2 platform, we found that the AS-3’UTR isoforms (3’UTR-1 and 3’UTR-2) of β-catenin mRNA are highly expressed in COAD, while the expression of unspliced UTR (3’UTR-3) isoform (annotated as the major transcript in the UCSC Genome Browser) is not significantly different between tumor and normal tissues (Fig. 2B). Notably, upregulation of 3′UTR-1 isoform was previously reported in liver hepatocellular carcinoma (LIHC), consistent with our GEPIA2 analysis (Supplementary Fig. S2B) (11). In contrast, expression of 3’UTR-2 was variable among LIHC patients, supporting its designation as an aberrant 3’UTR isoform of *CTNNB1*. Of note, we found consistent upregulation of aberrant 3’UTR-2 in CRC cell lines and COAD patients (Fig. 2B), suggesting its tissue-specific expression and potential tumorigenic roles in CRC.

**Fig. 2.**
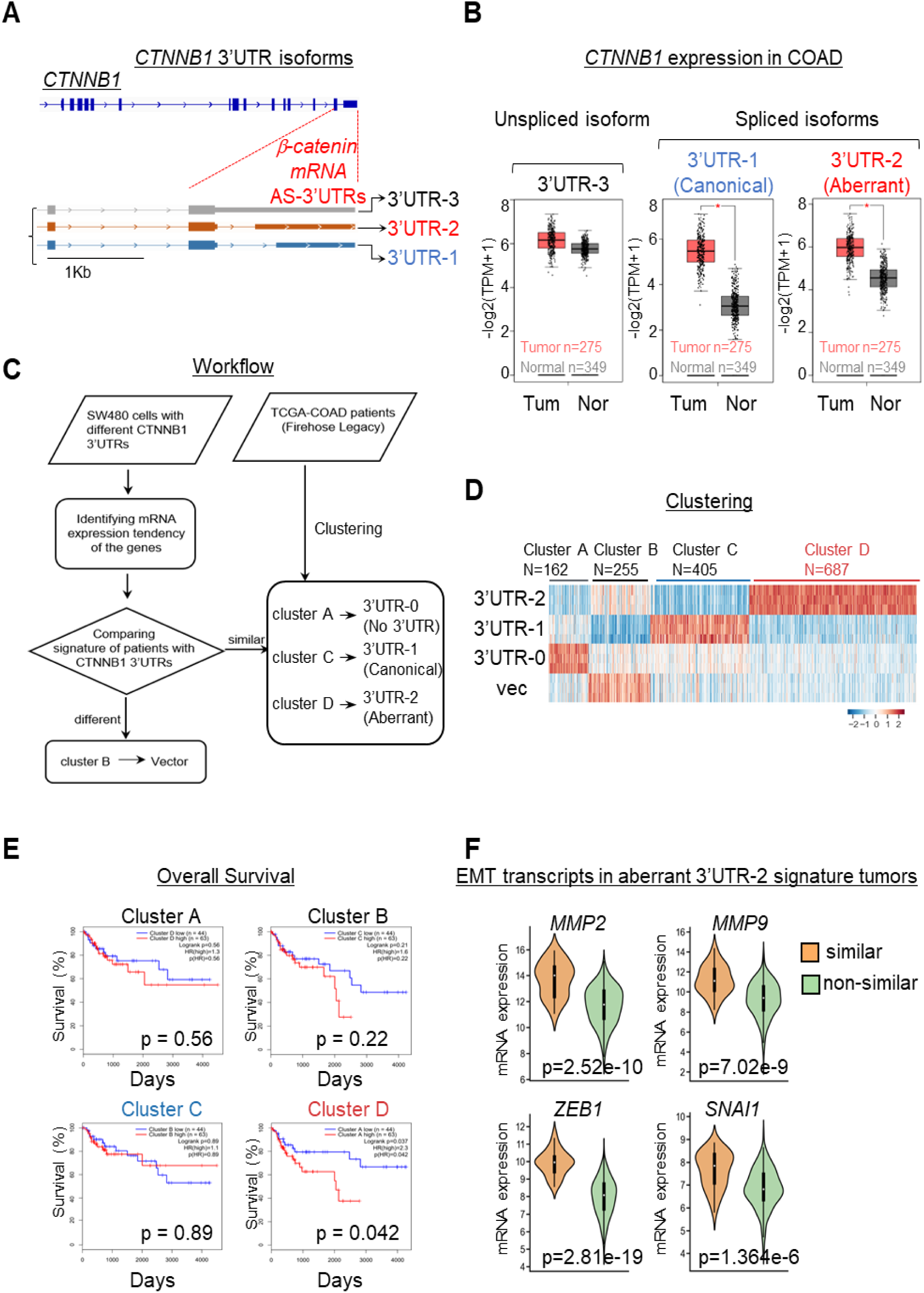
Aberrant *CTNNB1* 3′UTR isoforms associate with prognostic transcriptional signatures and poor survival in colon cancer. **(A)** Schematic representation of *CTNNB1* mRNA isoforms containing distinct 3′UTRs, including a normally spliced 3′UTR-1 (Canonical 3’UTR), an aberrantly spliced 3′UTR-2 (Aberrant 3’UTR) and an unspliced 3′UTR-3. **(B)** Expression profiles of *CTNNB1* 3′UTR isoforms in colon adenocarcinoma (COAD) patient samples and normal tissues analyzed using the GEPIA2 platform. **(C)** Workflow illustrating integration of RNA-sequencing datasets used for clustering analysis based on *CTNNB1* 3′UTR isoform expression. **(D)** Heatmap generated from K-means clustering of 1,509 differentially expressed genes identified from pairwise comparisons among four experimental conditions. **(E)** Kaplan–Meier analysis of overall survival in COAD patients stratified according to *CTNNB1* 3′UTR-associated transcriptional signatures. **(F)** Violin plots showing expression distributions of epithelial–mesenchymal transition (EMT) target genes across *CTNNB1* 3′UTR-defined clusters.

Upregulation of the aberrant 3’UTR-2 isoform prompted us to test how AS-3’UTR isoforms influence clinical outcomes in CRC and COAD patients. To evaluate the transcriptional signature by β-catenin mRNA AS-3’UTRs, we established the CRC cell lines stably expressing FLAG-β-catenin with distinct AS-3’UTR isoforms (Supplementary Fig. S2C and S2D). Integrated RNA-seq revealed differentially expressed genes (DEGs) stratified into four clusters (A-D) according to their expression signatures of different β-catenin AS-3’UTRs (Fig. 2C and 2D). Cluster D corresponded to the transcriptional profile of cells expressing β-catenin mRNA with aberrant 3’UTR-2 isoform (Fig. 2D). Notably, overall survival was significantly reduced in the Cluster D signature COAD patients, while clusters A-C demonstrated no significant prognostic differences (Fig. 2E). Importantly, COAD patients with 3’UTR-2 transcriptional signature resemblance (Cluster D) exhibited high expression of key epithelial-to-mesenchymal transition (EMT) and metastasis-related genes, including *MMP2, MMP9, ZEB1*, and *SNAI1,* as in the 3’UTR-2 expressing tumors (Fig. 2F). These imply a functional relevance of an aberrant 3’UTR-2 in promoting EMT-associated invasive phenotypes in COAD. Collectively, aberrant 3′UTR-2 isoform may drive transcriptional programs associated with aggressive tumor behavior.

### An aberrant 3′UTR of β-catenin mRNA reprograms transcriptional programs

To systematically investigate the functional impact of AS-3’UTR isoforms in CRC, we established stable cell lines of β-catenin CRISPR knockout followed by FLAG-β-catenin rescue. β-catenin protein was reconstructed with distinct AS-3’UTR mRNA isoforms: 3’UTR-0 (β-catenin mRNA coding region, but no 3’UTR), 3’UTR-1 (coding region and 3’UTR with Exon 16B), and 3’UTR-2 (coding region and 3’UTR with Exon 16A and 16B, referred to as aberrant 3’UTR) (Fig. 2A and Supplementary Fig. S3A-S3B). RNA-seq analysis of stable cell lines identified 1,026 genes downregulated upon β-catenin CRISPR knockout, of which 144 were significantly restored in cells expressing β-catenin-3’UTR-2, compared to 25 and 47 genes in the 3’UTR-0 and 3’UTR-1 groups, respectively (p<0.01; Fig. 3A and Supplementary Table 2A-S2E). Wnt/β-catenin target genes, including proliferative genes, were downregulated upon knockout and rescued by all β-catenin genes, suggesting that FLAG-β-catenin proteins act as transcription factors regardless of AS-3’UTRs of mRNA (Fig. 3B and Supplementary Fig. S3C-S3E). In contrast, distinct gene sets were activated depending on AS-3’UTRs: stem cell differentiation genes were upregulated by 3’UTR-1 and cell morphogenesis-related signatures were altered by 3’UTR-2 (Fig. 3B). These results indicate distinct AS-3’UTR isoforms differentially regulate β-catenin target genes, thereby reprogramming the transcriptional landscape.

**Fig. 3.**
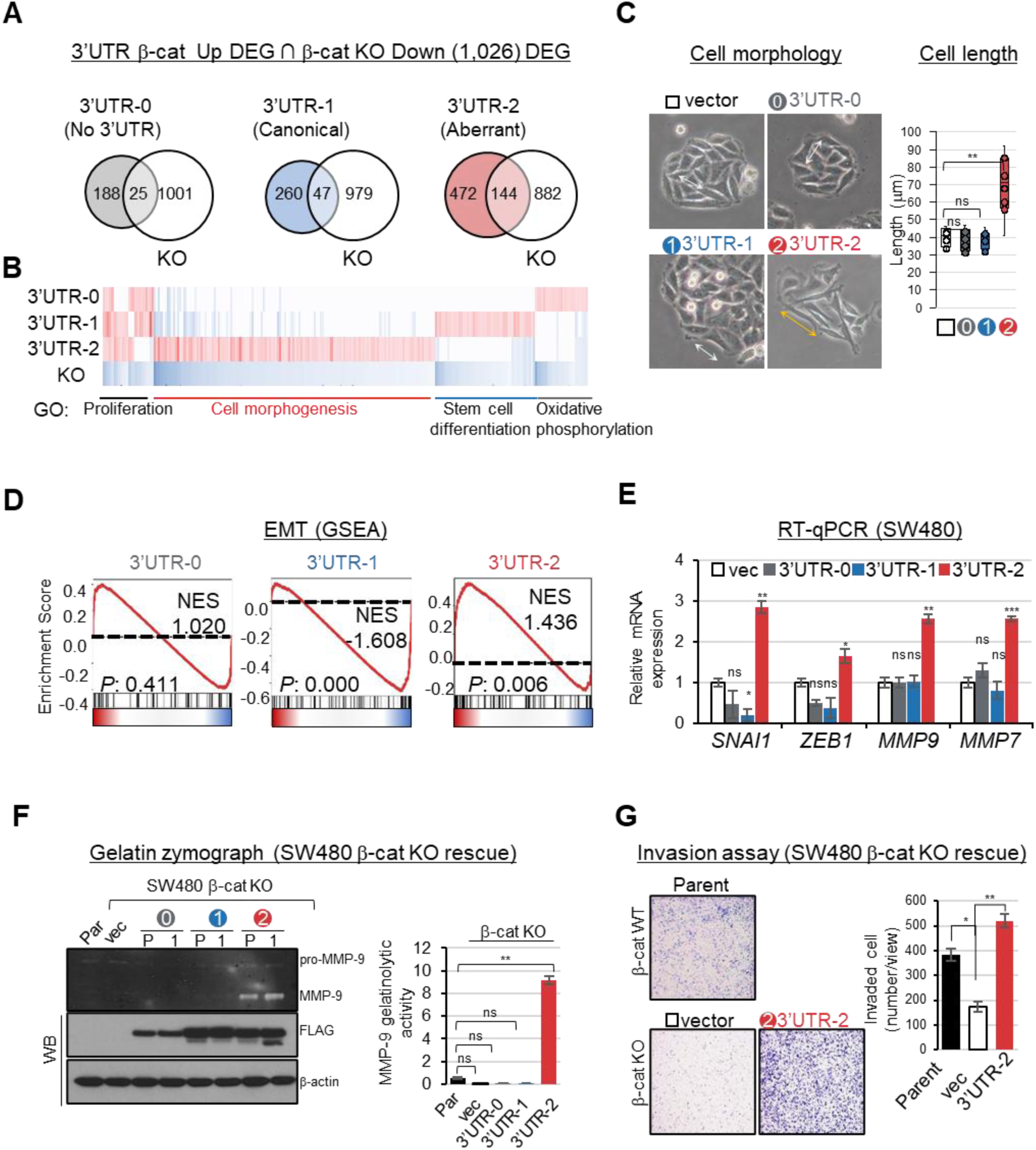
Aberrant 3′UTRs promote EMT-associated transcriptional and invasive phenotypes. **(A)** Venn diagram showing the number of differentially expressed genes identified across RNA-sequencing datasets comparing β-catenin 3′UTR isoform-expressing cells. **(B)** Heatmap showing differential gene expression patterns among cells expressing β-catenin 3′UTR variants. **(C)** Representative phase-contrast images showing morphological changes in SW480 cells expressing FLAG–β-catenin constructs containing distinct 3′UTRs (left). Quantification of cell size and morphological descriptors is shown (N > 20 cells per condition). **(D)** Gene set enrichment analysis (GSEA) of RNA-seq data from SW480 cells expressing β-catenin 3′UTR variants, highlighting enrichment of epithelial–mesenchymal transition (EMT)-associated gene sets. **(E)** Validation of selected transcript changes by RT–qPCR. **(F)** Gelatin zymography showing matrix metalloproteinase-9 (MMP-9) activity in conditioned medium from SW480 cells expressing vector control (vec), 3’UTR-0, 3’UTR-1, and 3’UTR-2 β-catenin constructs. **(G)** Matrigel invasion assay of SW480 cells expressing β-catenin 3′UTR variants or vector control. Invaded cells on the lower surface of transwell inserts were quantified at 24, 48, and 72 h (n = 3).

As cell morphogenesis-related genes were upregulated, we observed EMT-like morphological changes and increased cell size in UTR-2 stable cells (Fig. 3C). Gene set enrichment analysis (GSEA) revealed significant enrichment of EMT-related genes in the 3’UTR-2 cells, accompanied by increased expression of EMT-associated transcription factors (*SNAI1* and *ZEB1*) both in SW480 and HT-29 stable cells (Fig. 3D and Supplementary Fig. S4A and Supplementary Table 3). Expression of metalloproteinase genes (*MMP7* and *MMP9*) for extracellular matrix (ECM) remodeling was elevated, and gelatin zymography activity of proMMP-9 was enhanced in aberrantly spliced 3’UTR-2 cells (Fig. 3E and 3F). Finally, transwell migration and invasion assays demonstrated increased invasiveness in β-catenin-3’UTR-2 expressing cells (Fig. 3G and Supplementary Fig. S4B-S4C). Collectively, these findings indicate that β-catenin expressed from the aberrant 3′UTR-2 drives an EMT-prone transcriptional and phenotypic states. These transcriptional changes suggested that AS-3′UTRs may alter β-catenin localization through RNA-based mechanisms.

### An aberrant β-catenin 3′UTR forms cytoplasmic RNA condensates through interaction with FUS

3’UTRs are known to regulate mRNA stability, translational efficiency, and subcellular localization. Since mRNA stability and translation efficiency were not significantly different in cells with AS-3’UTRs of β-catenin mRNA (Supplementary Fig. S5A-S5B), we next investigated whether the aberrant 3’UTR alters subcellular localization and molecular interactions of β-catenin mRNA. To address this, we first examined the distribution of endogenous β-catenin mRNA coding sequences using RNAscope and single molecule fluorescence in situ hybridization (smFISH) in HCT116 and SW480 cells (Fig. 4A and Supplementary Fig. S5C-S5D). The β-catenin mRNA was detected both in the nucleus and cytoplasm, indicating broad subcellular distribution of β-catenin transcripts in CRC cells (Fig. 4B and Supplementary Fig. S5D). To specifically localize individual 3’UTR isoforms, we designed smFISH probes targeting specific splicing junctions of 3’UTR isoforms (Exon15/16B for 3’UTR-1, Exon15/16A for 3’UTR-2, Exon15/In15 for 3’UTR-3). The unspliced isoform (3′UTR-3), which retains an intron and corresponds to the annotated major transcript, was predominantly retained in the nucleus. In contrast, the spliced isoforms (3’UTR-1 and 3’UTR-2) were exported to the cytoplasm (Figs. 4C, 4D and Supplementary Fig. S5E). Remarkably, aberrant 3′UTR-2 isoform was predominantly localized in the cytoplasm, where it formed discrete RNA condensates, while 3’UTR-1 was broadly distributed throughout the nucleus and the cytoplasm (Fig. 4C, 4D and Supplementary Fig. S5F). Because RNA condensates are often mediated by RNA–protein interactions, we hypothesized that specific RBPs recognize the aberrant 3′UTR-2 isoform to drive condensate formation.

**Fig. 4.**
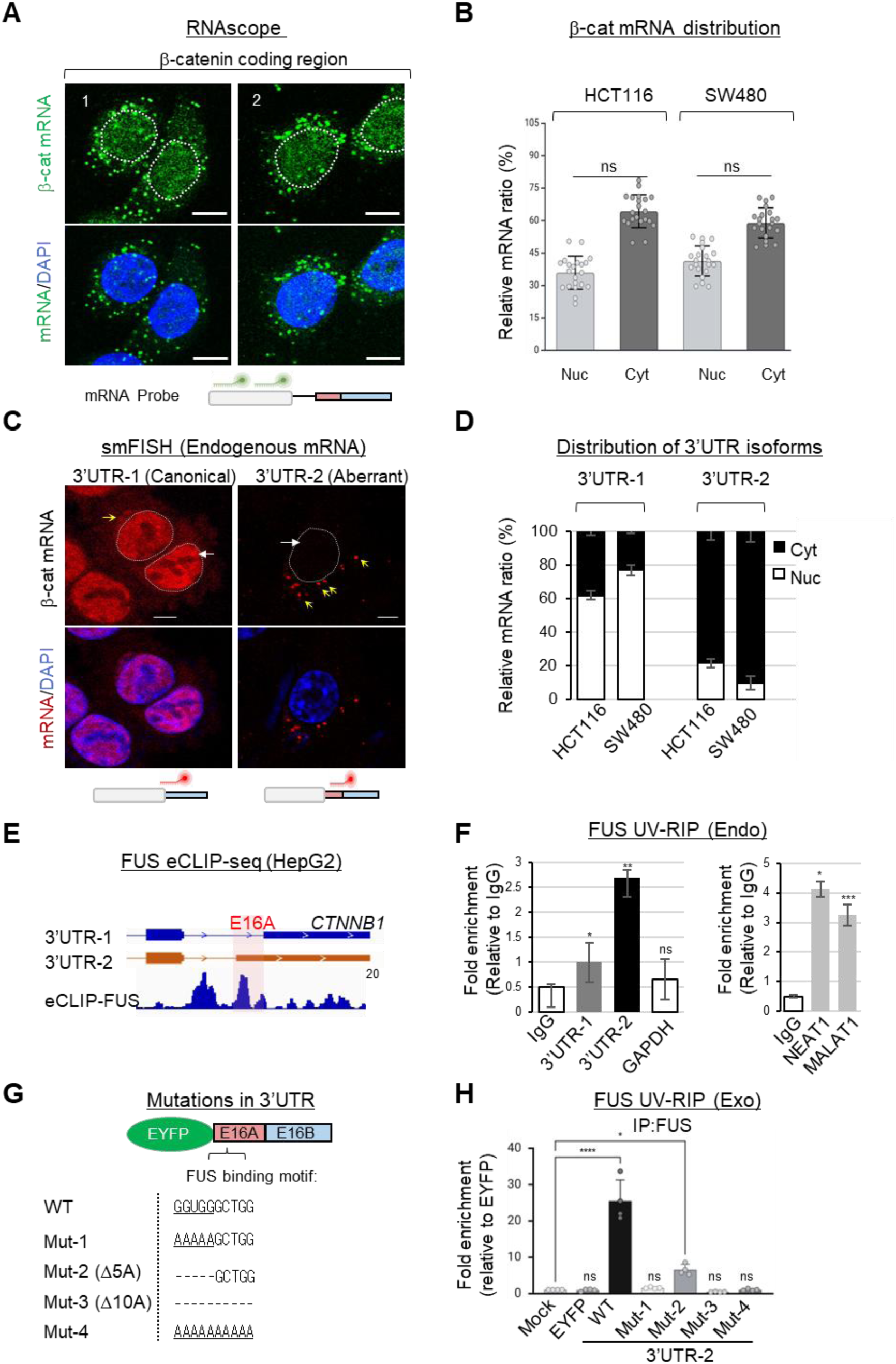
FUS binds an aberrant 3′UTR exon and promotes cytoplasmic β-catenin mRNA condensates. **(A)** RNAscope images showing subcellular localization of total β-catenin mRNA in SW480 cells. **(B)** Quantification of the relative distribution of β-catenin mRNA between cytoplasmic and nuclear compartments in HCT116 and SW480 cells. **(C)** Single-molecule fluorescence in situ hybridization (smFISH) images detecting β-catenin 3′UTR-1 and 3′UTR-2 isoforms using junction-specific probes in SW480 cells. Yellow arrows indicate cytoplasmic mRNA granules, and white arrows indicate nuclear-localized transcripts. **(D)** Quantification of the relative distribution of β-catenin 3′UTR isoforms between cytoplasmic and nuclear compartments in SW480 cells. **(E)** FUS enhanced crosslinking and immunoprecipitation sequencing (eCLIP-seq) tracks aligned to β-catenin transcripts showing enriched binding at alternative exon 16A. Predicted FUS-binding motifs identified by RBPmap are indicated. **(F)** UV-RNA immunoprecipitation (UV-RIP) analysis showing FUS association with β-catenin 3′UTRs compared with GAPDH (negative control) and NEAT1 and MALAT1 (positive controls), respectively. **(G)** Schematic representation of reporter constructs containing wild-type or mutant FUS-binding motifs within the alternative exon 16A. **(H)** RNA-IP–qPCR analysis showing FUS binding to alternative exon 16A using reporter constructs described in **G** following UV crosslinking in SW480 cells.

Next, we sought to identify RNA binding proteins (RBPs) for exclusive cytoplasmic RNA localization of aberrant 3’UTR-2 (Supplementary Fig. S6A and Supplementary Table 4). Candidate RBPs recognizing the alternative exon (Exon 16A) unique to 3’UTR-2 were predicted using RBPmap and analyzed with publicly available eCLIP-seq datasets (GSE177328), nominating FUS protein as a strong candidate through enriched GGUGG motifs in Exon 16A but not in Exon 16B (Fig. 4E and Supplementary Fig. S6A-6E) (57–59). To validate the direct interaction between FUS and β-catenin mRNA, we performed UV-RNA Immunoprecipitation (UV-RIP) using anti-FUS antibodies and analyzed bound β-catenin mRNA, showing preferential FUS binding with 3’UTR-2 in comparison to 3’UTR-1 (Fig. 4F). *NEAT1* and *MALAT1* served as a positive control, while *GAPDH* was used as negative controls, similar with previously reported FUS eCLIP-seq datasets (Fig. 4F and Supplementary Fig. S6F). FUS binding preference was also examined by UV-RIP with 3’UTR reporters. Mutation analysis of 3’UTR reporters demonstrated that the mutations of FUS-binding motif in Exon 16A significantly reduced FUS association (Figs. 4G, 4H and Supplementary Fig. S6G). These results demonstrate that FUS specifically binds to the Exon 16A in aberrant 3’UTR-2, supporting a role for FUS in isoform-specific RNA localization and condensate formation.

### FUS regulates localization and translation of β-catenin mRNA

We then directly tested whether FUS modulates the subcellular localization of β-catenin 3’UTR-2 mRNA, by immunofluorescence and smFISH in CRC cells. FUS was predominantly nuclear; however, cytoplasmic FUS strongly correlated with the presence of 3′UTR-2 mRNA granules (R^2^ = 0.87) (Figs. 5A and 5B). Consistently, in SW480 cells, robust colocalization between cytoplasmic FUS and 3’UTR-2 mRNA granules were observed (Fig. 5A). To determine whether FUS is required for granule formation, we knocked down FUS protein expression. Loss of FUS markedly reduced both the number and size of 3’UTR-2 mRNA-containing granules, as well as their colocalization with FUS (Figs. 5C and 5D).

**Fig. 5.**
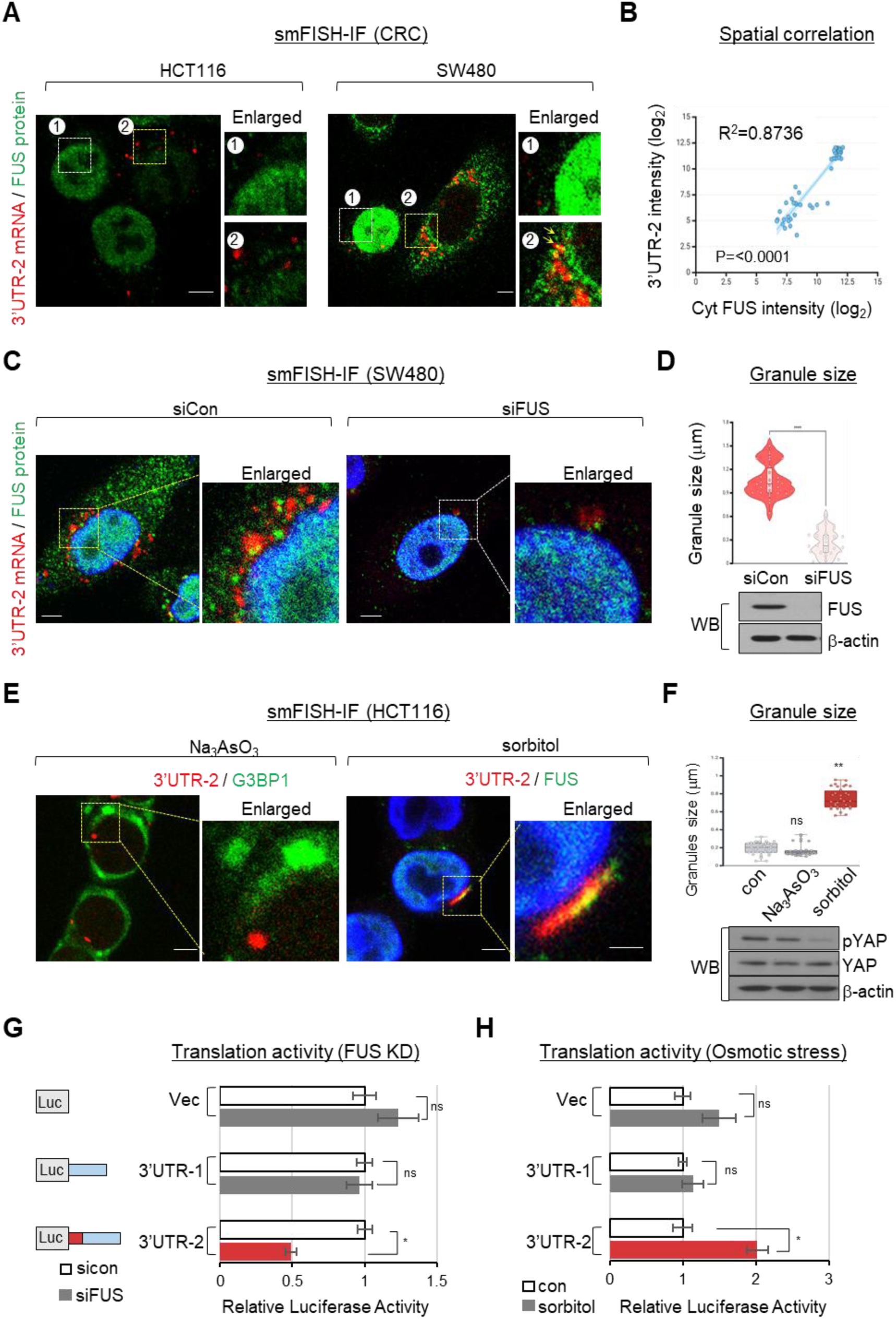
FUS regulates cytoplasmic condensate formation and translation of the β-catenin mRNA harboring an aberrant 3′UTR. **(A)** smFISH and immunofluorescence images showing colocalization of β-catenin 3′UTR-2 mRNA with FUS in HCT116 and SW480 cells. **(B)** Correlation analysis of β-catenin 3′UTR-2 mRNA granule intensity and cytoplasmic FUS signal (R² = 0.8736, *P* ≤ 0.0001). **(C)** Dual smFISH and immunofluorescence showing the effect of FUS knockdown on β-catenin 3′UTR-2 mRNA localization in SW480 cells. **(D)** Violin plot quantifying cytoplasmic β-catenin 3′UTR-2 mRNA granule size following FUS depletion (top). Immunoblot validating FUS knockdown efficiency (bottom). **(E)** smFISH and immunofluorescence of β-catenin 3′UTR-2 mRNA with G3BP1 in sodium arsenite (Na₃AsO₃)-treated HCT116 cells, and with FUS in sorbitol-treated cells. **(F)** Quantification of β-catenin 3′UTR-2 mRNA and FUS colocalization under indicated stress conditions (top). Immunoblot showing phosphorylated YAP (p-YAP) and total YAP as stress response controls (bottom). β-actin was used as a loading control. **(G and H)** Luciferase reporter activity of β-catenin 3′UTR constructs (3′UTR-1 and 3′UTR-2) following FUS depletion (siFUS) in SW480 cells **(G)** and after sorbitol treatment in HCT116 cells **(H)**.

We next examined the identity of these granules using stress granule marker G3BP1 following sodium arsenite-induced oxidative stress. 3’UTR-1 was colocalized with G3BP1, indicating stress granule localization of β-catenin mRNA; in contrast, 3’UTR-2 mRNA granules did not co-localize with G3BP1, suggesting that 3’UTR-2 RNA condensates are distinct from canonical stress granules (Fig. 5E and Supplementary Fig. S7A-S7B). To further assess stress responsiveness, hypertonic stress was induced by sorbitol, which triggers nuclear export and cytoplasmic accumulation of FUS protein (60–62). Under these conditions, colocalization between FUS and 3’UTR-2 mRNA granule was significantly enhanced, whereas the localization of 3’UTR-1 mRNA remained unchanged (Fig. 5E, and Supplementary Fig. S7C). Sorbitol treatment did not alter overall β-catenin mRNA levels, as the experimental conditions validated by YAP dephosphorylation, positive control for sorbitol-induced stress response (Fig. 5F and Supplementary Fig. S7D).

Finally, luciferase reporter assays demonstrated that the 3′UTR-2 isoform enhances translational efficiency in an FUS-dependent manner. Depletion of FUS significantly reduced translation activity of the 3’UTR-2 reporter but not 3’UTR-1 reporter (Fig. 5G). Similarly, sorbitol-induced stress increased translation of reporters containing AS-3′UTRs (Fig. 5H). Because the β-catenin coding sequence was replaced with luciferase, these results indicate that FUS-dependent regulation of translation is mediated through the AS-3′UTR. Together, these results demonstrate that FUS promotes formation of cytoplasmic β-catenin mRNA condensates that enhance translation under stress-responsive conditions.

### β-catenin protein localization is directed by the alternative exon within the aberrant 3’UTR

The distinct subcellular localization of β-catenin AS-3’UTRs promoted us to ask whether this regulatory activity of AS-3’UTRs extends to the localization of β-catenin protein. We re-used *CTNNB1* knockout cells reconstituted with FLAG-β-catenin carrying distinct AS-3′UTR (3′UTR-0, -1, and -2) (Supplementary Fig. S3, Fig. 3). Immunofluorescence and cell fractionation analyses revealed isoform-dependent localization of β-catenin protein: β-catenin expressed from 3’UTR-0 and 3’UTR-1 was predominantly localized in the nucleus and plasma membrane, whereas β-catenin derived from 3’UTR-2 was primarily localized in the cytoplasm (Figs. 6A-6C).

**Fig. 6.**
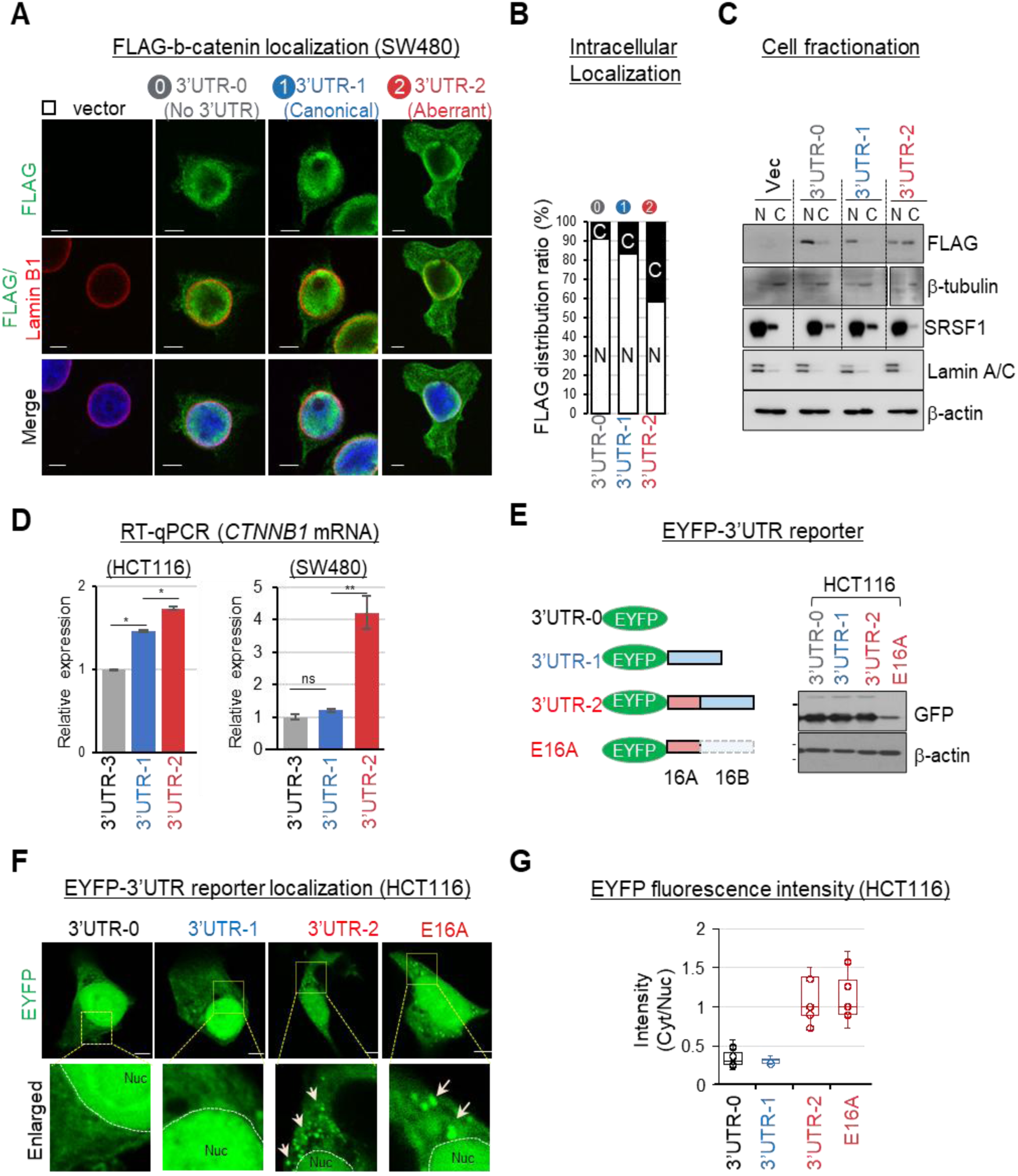
An aberrant 3′UTR retargets subcellular localization of β-catenin protein. **(A)** Immunofluorescence of FLAG-tagged β-catenin (green) and Lamin B1 (red) in SW480 cells expressing FLAG–β-catenin constructs harboring distinct 3′UTRs following endogenous β-catenin knockout. **(B)** Quantification of FLAG–β-catenin subcellular distribution ratios. **(C)** Immunoblot analysis of cytoplasmic and nuclear fractions from SW480 cells expressing β-catenin 3′UTR variants. **(D)** Quantitative RT–PCR analysis of β-catenin 3′UTR isoforms in HCT116 and SW480 cells. **(E)** EYFP–3′UTR reporter constructs used to investigate functional domains within the β-catenin 3′UTR. Immunoblot showing EYFP expression in HCT116 cells. **(F)** Immunofluorescence images of EYFP–3′UTR constructs in SW480 cells. **(G)** Quantification of cytoplasmic-to-nuclear distribution ratios of EYFP signal in SW480 and HCT116 cells.

Because a distinct localization pattern was observed for the 3’UTR-2, we next tested whether Exon 16A within 3’UTR-2 is sufficient to direct subcellular distribution of β-catenin protein. Expression of 3’UTR-1 and 3’UTR-2 isoforms of *CTNNB1* mRNA was analyzed using splice junction-specific primers in CRC cells, revealing similar basal levels of AS-3’UTRs in HCT116 cells (Fig. 6D). To determine whether specific 3’UTR elements control protein localization, we expressed EYFP reporter constructs containing full-length AS-3’UTRs or individual fragments (3’UTR-1, 3’UTR-2 or the alternative exon within 3’UTR-2 [Exon 16A]) in HCT116 cells (Fig. 6E). EYFP expression levels were comparable between 3’UTR-1 and 3’UTR-2 constructs, whereas expression from the Exon 16A-only construct was slightly reduced (Fig. 6E and Supplementary Fig. S8A). Strikingly, Exon 16A specifically promoted cytoplasmic localization of EYFP (Figs. 6F and 6G). Consistent with this observation, Exon 16A alone was sufficient to direct EYFP to the cytoplasm in SW480 cells (Supplementary Fig. S8B).

Together, our data support an unexpected role of AS-3’UTR isoforms, particularly the alternative Exon 16A, in directing protein localization. Because subcellular localization of β-catenin mRNA was regulated by the aberrant 3’UTR, these results indicate that protein localization is modulated by AS-3’UTRs through isoform-specific RNA localization, linking 3’UTR splicing to spatial control of protein function. These findings raised the possibility that altered cytoplasmic localization of β-catenin encoded by 3′UTR-2 may impair its recruitment to membrane-associated adhesion complexes.

### β-catenin-E-cadherin complex formation is regulated by AS-3’UTR of β-catenin mRNA

Given the multifaceted roles of β-catenin protein as both a WNT-activated nuclear transcription factor and a membrane-bound adherens junction (AJ) protein (31,32), we hypothesized that distinct AS-3’UTR isoforms may contribute to its functional diversification by guiding the localization and protein complex formation. While multiple *CTNNB1* 3’UTR isoforms have been reported, the functional significance of these isoforms in regulating the dual roles of β-catenin has remained unclear. Co-immunoprecipitation demonstrated that FLAG-β-catenin proteins derived from all 3’UTR variants could bind TCF4, and TOP/FOP luciferase reporter assays confirmed comparable transcriptional activity among the isoforms, consistent with preservation of β-catenin nuclear transcriptional function across isoforms (Fig. 7A and Supplementary Fig. S9A). Consistently, canonical WNT targets such as *CCND1, AXIN2, and c-MYC* were upregulated in all isoform-expressing cells (Fig. 7B). These results indicate that the transcriptional activity of β-catenin is primarily determined by its coding region independent of 3’UTR usage.

**Fig. 7.**
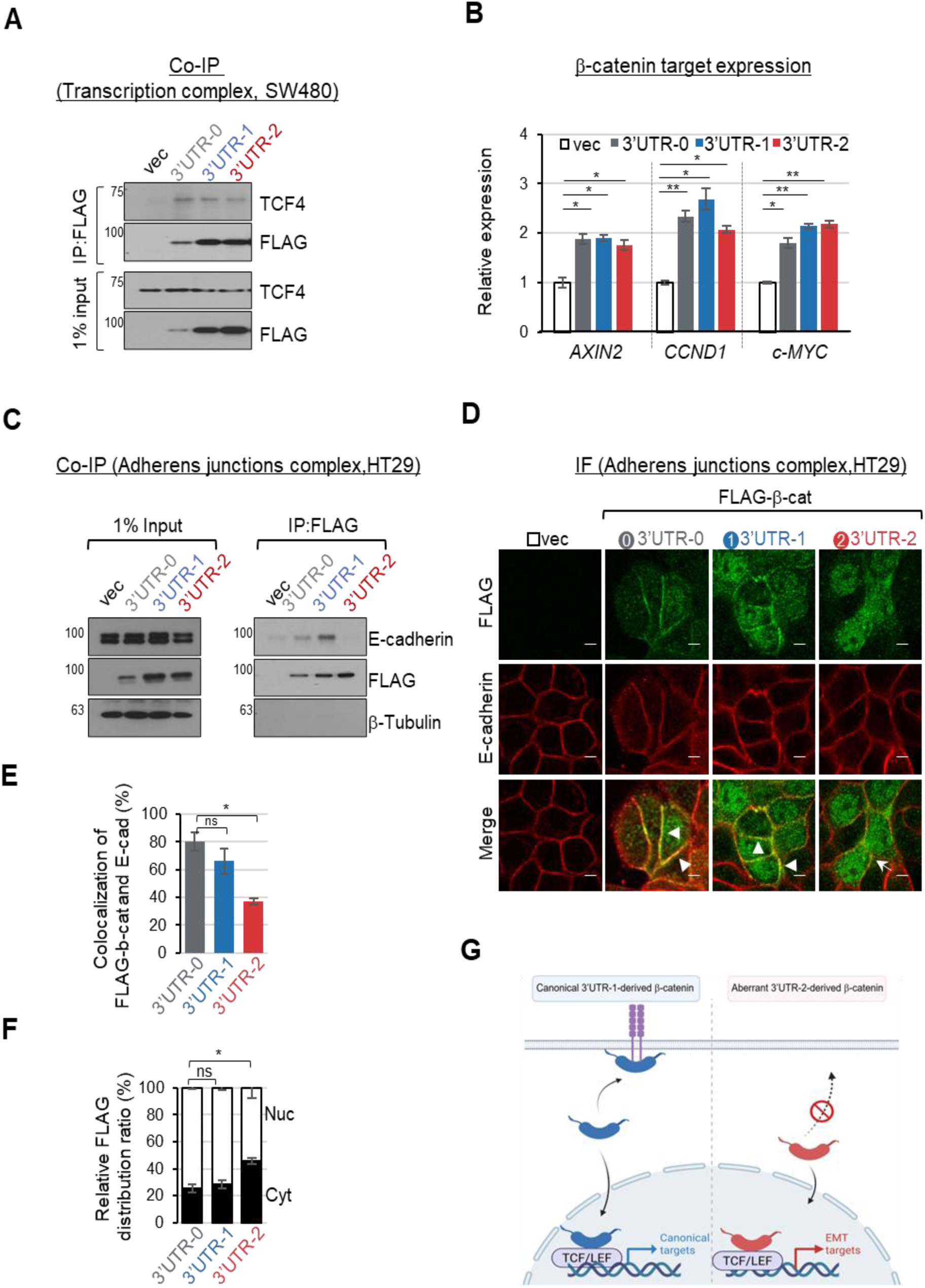
An aberrant 3′UTR disrupts β-catenin adhesion complex formation. **(A)** FLAG immunoprecipitation from SW480 cells expressing FLAG–β-catenin constructs harboring distinct 3′UTR variants for detection of endogenous TCF4 (nuclear interaction control). **(B)** Quantitative RT–PCR analysis of β-catenin/TCF4 target gene expression in SW480 cells expressing FLAG–β-catenin constructs harboring distinct 3′UTR variants. **(C)** FLAG immunoprecipitation from HT-29 cells expressing FLAG–β-catenin constructs harboring distinct 3′UTR variants for detection of endogenous E-cadherin. **(D)** Immunofluorescence staining for FLAG–β-catenin and E-cadherin (an adherens junction marker; red) in HT-29 cells expressing FLAG–β-catenin constructs harboring distinct 3′UTR variants. **(E)** Quantification of colocalization between FLAG–β-catenin and E-cadherin in cells expressing FLAG–β-catenin constructs harboring distinct 3′UTR variants. **(F)** Quantification of the subcellular distribution of FLAG–β-catenin proteins in cells expressing FLAG–β-catenin constructs harboring distinct 3′UTR variants. **(G)** Proposed model illustrating that β-catenin derived from the normally spliced 3′UTR-1 isoform interacts with both TCF4 and E-cadherin, whereas β-catenin derived from the aberrant 3′UTR-2 isoform reduced interaction with E-cadherin at adherens junctions.

Because AJ assembly is essential for maintaining epithelial integrity and β-catenin plays an essential role in cell adhesion complexes, we next tested whether AS-3′UTR isoforms differentially regulate cell–cell adhesion complex formation and invasive phenotypes as observed in 3′UTR-2–expressing cells (Supplementary Fig. S4B-4C). We utilized HT-29 cells, which exhibit robust cell-cell adhesion mediated by E-cadherin. Efficient interaction with E-cadherin was observed for β-catenin proteins derived from 3’UTR-0 and 3’UTR-1 (Fig. 7C and Supplementary Fig. S9B). In contrast, β-catenin protein encoded by the aberrant 3’UTR-2 isoform failed to interact with E-cadherin (Fig. 7C). Immunofluorescence analysis further confirmed that β-catenin derived from 3’UTR-0 and 3’UTR-1 localized at the plasma membrane, where it colocalized with E-cadherin at AJs in HT-29 cells (Figs. 7D and 7E). By contrast, FLAG-β-catenin encoded by the 3’UTR-2 isoform was predominantly localized in the cytoplasm and excluded from AJs (Figs. 7D-7F).

Together, our data demonstrate that while the transcriptional activity of β-catenin is largely independent of 3’UTR usage, its ability to participate in AJ complexes is critically regulated by 3’UTR isoform identity (Fig. 7G). These results provide a mechanistic explanation for how β-catenin transcriptional activity is uncoupled from adhesion function, and links cancer-associated 3′UTR dys-splicing to the disruption of AJs to invasive phenotypes in colorectal cancer.

## Discussion

Our study uncovers a previously unrecognized regulatory layer in which 3′UTR alternative splicing, rather than coding sequence change, determines the subcellular fate of β-catenin protein. Although AS in coding regions has long been recognized as a primary driver of proteomic diversity, our findings demonstrate that AS-3′UTRs-driven transcriptome diversity can reprogram protein behavior without changing an amino acid sequence. We show that AS-3′UTRs are widespread across cancer-relevant transcripts, supporting their generality. Using β-catenin mRNA as a model, we establish that an aberrantly spliced 3′UTR isoform drives FUS-dependent cytoplasmic mRNA condensate formation, prevents β-catenin incorporation into E-cadherin-based adherens junctions, and induces EMT-associated transcriptional programs, collectively linking aberrant 3′UTR splicing to cancer aggressiveness.

Several cancer-relevant transcripts harbor prognostically significant AS-3’UTRs. For example, *TCF3* (also known as E2A) exhibits two distinct 3’UTR isoforms with differential association in colorectal adenocarcinoma (COAD): the unspliced isoform is enriched in tumors, whereas the spliced isoform is reduced compared with normal tissues (63–65). *EIF3B*, a core component of the translation initiation factor complex, has also been implicated in tumorigenesis and poor prognosis across multiple cancers (66,67). Importantly, our findings indicate that specific alternatively spliced 3’ UTR isoforms can modulate adverse clinical impact, suggesting their potential as candidate therapeutic targets. These observations support the concept that AS-3’UTRs provide an underappreciated regulatory layer, enabling non-coding regions of mRNAs to diversify protein function and clinical outcome without altering amino acid sequence.

mRNA localization regulates localized translation and protein targeting, often influencing protein fate through co-translational interactions (12,68–70). Additionally, mRNA localization combined with translation rate can determine protein targeting to multiple subcellular destinations by modulating interactions with binding partners (71). Isoform-specific mRNA localization has emerged as an important mechanism for spatial regulation of gene expression. Emerging evidence indicates that mRNA isoforms influence protein multifunctionality by directing how proteins are synthesized and function within cells. Multiple 3’UTR isoforms can differentially regulate transcript localization and translation without altering the encoded amino acid sequence. For example, alternative 3′UTR isoforms of *CDC42* and *Pumilio* exhibit distinct localization patterns, with longer isoforms retained near the cell body and shorter isoforms transported to distal compartments to regulate local protein synthesis (10,72). Recent studies have shown that alternative 3’UTRs regulate RNA surveillance and transcript stability pathways such as nonsense-mediated decay (NMD) (79). In parallel, pan-cancer analyses have reported widespread alternative splicing of the *CTNNB1* 3′UTR in hepatocellular carcinoma and other cancer types (11), supporting the broader relevance of alternative 3’UTR regulation across malignancies. However, these studies did not address how alternative 3’UTR isoforms contribute to functional diversification and subcellular localization of their associated proteins. Our findings suggest that AS-3′UTR isoforms encode spatial regulatory information that directs β-catenin localization and activity, thereby providing a mechanistic link between RNA processing and spatial regulation of b-catenin function.

Accumulating evidence highlights spatial control of β-catenin protein expression and function. In neuronal cells, β-catenin mRNA is transported to axons and locally translated at presynaptic boutons, enabling localized accumulation of β-catenin (73). This process is regulated by cytoplasmic polyadenylation element-binding protein-1 (CPEB1) (74). These studies illustrate that spatial regulation of mRNA translation can control protein function within specific cellular compartments. Consistent with this principle, our results demonstrate that β-catenin AS-3’UTR mRNA isoforms contribute to isoform–dependent spatial regulation of protein localization in cancer cells. The β-catenin protein translated from the aberrant 3′UTR-2 exhibited altered intracellular localization.

β-catenin protein derived from 3′UTR-2 exhibited reduced membrane association and disrupted interaction with E-cadherin. Impaired formation of adhesion complexes is consistent with observed changes in β-catenin protein subcellular distribution. This spatial regulation appears to be linked to cytoplasmic mRNA condensates through interaction with FUS, thereby contributing to altered localization of the 3′UTR-2 isoform. These findings suggest that alternative 3′UTR usage can modulate the subcellular distribution of β-catenin protein through mRNA localization, thereby altering protein interaction partners and functional outputs in cancer cells. Together, we propose a model in which aberrant 3′UTR splicing reprograms protein targeting through mRNA condensate–mediated spatial regulation.

Although β-catenin is a key driver of tumor initiation, its role in metastatic progression remains incompletely understood (33,38,75). β-catenin functions as a membrane-associated component of adherens junctions through interaction with E-cadherin, thereby maintaining cell–cell adhesion and epithelial integrity. Dissociation of β-catenin from the E-cadherin complex leads to cytoplasmic accumulation and nuclear translocation, where β-catenin activates EMT-associated gene expression programs (75,76). Indeed, activation of Wnt/β-catenin signaling induces EMT-associated transcription factors such as *SNAI1* (*SNAIL*) and *SLUG* (77,78). Despite extensive investigation on β-catenin signaling, mechanisms governing its spatial regulation remain incompletely defined. Our data support a functional role of AS-3’UTRs of β-catenin mRNA as critical determinants of its intracellular localization and function, particularly in the context of adhesion complex formation. We demonstrate that alternative 3′UTR isoforms of β-catenin mRNA led to distinct subcellular localization patterns and influence downstream functional outcomes. β-catenin protein translated from the 3′UTR-2 was associated with enrichment of EMT-related transcriptional programs, along with increased migratory and invasive capacities of colorectal cancer cells. Clinical analysis further supported the relevance of this mechanism, as colorectal cancer (COAD) patients exhibiting high expression of β-catenin 3′UTR-2-associated gene signatures showed significantly poorer overall survival, implicating the aberrant 3′UTR–2 mRNA isoform as a potential contributor to colorectal cancer progression.

Collectively, our findings support a model in which aberrant AS-3′UTRs function as spatial regulatory elements that reprogram protein localization and cellular behavior through RNA condensate-mediated mechanisms. Further studies will be necessary to determine how specific AS-3′UTR sequences or structures promote FUS-dependent condensate formation, potentially leading to distinct cytoplasmic functions of β-catenin in cancer cells.

## Abbreviation

3’UTR: 3’ Untranslated Region
AS: Alternative Splicing
RBP: RNA Binding Protein
EMT: Epithelial-to-Mesenchymal Transition
AJ: Adherens Junction

