## Supplementary information for "Aberrant 3′UTR splicing drives FUS-dependent mRNA condensates and prevents β-catenin from adherens junctions to promote cancer aggressiveness"

**Table information**

Table 1 List of alternative splicing 3'UTR (AS-3'UTR)

Table 2 List of differentially expressed genes in RNA-seq

Table 3 List of EMT gene in GSEA results

Table 4 List of RNA binding proteins by RBPmap

Table 5 List of materials used in the study

### Supplementary Figures

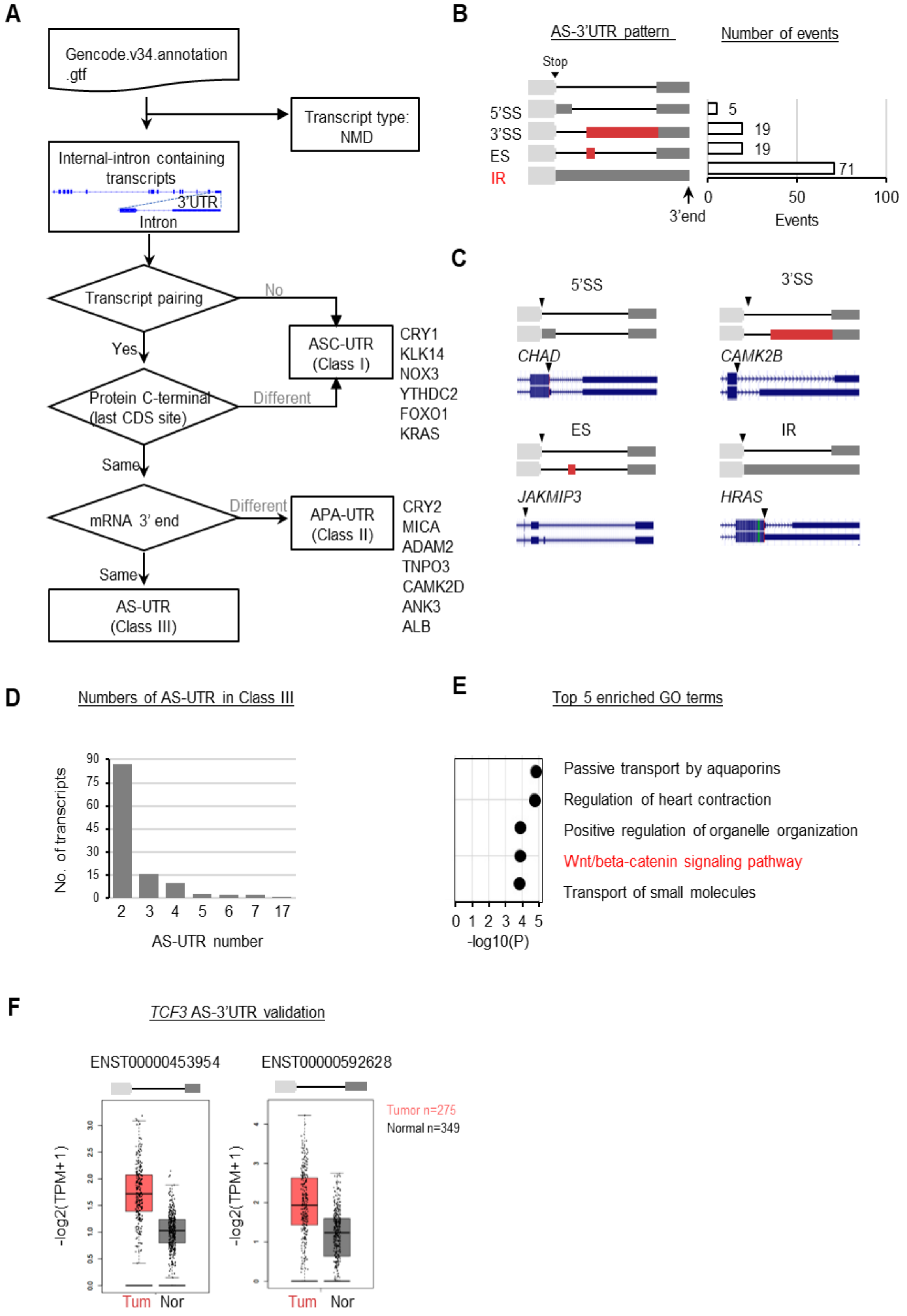

**Supplementary Figure 1. Characterization of alternative splicing in 3’UTRs**

**(A)** Workflow for identifying alternative splicing (AS) events in the 3’UTRs. **(B)** Schematic of five differential patterns within 3’UTRs and the number of genes associated with each pattern. **(C)** Representative genome browser views of four example genes showing distinct AS-3’UTR splicing patterns. **(D)** Distribution of the number of AS-3’UTR isoforms per transcript in Class III. **(E)** Gene Ontology (GO) enrichment analysis showing the top 10 biological processes associated with Class III AS-3’UTRs. **(F)** Kaplan–Meier analysis showing overall survival in COAD patients stratified by *TCF3* AS-3′UTR isoform expression.

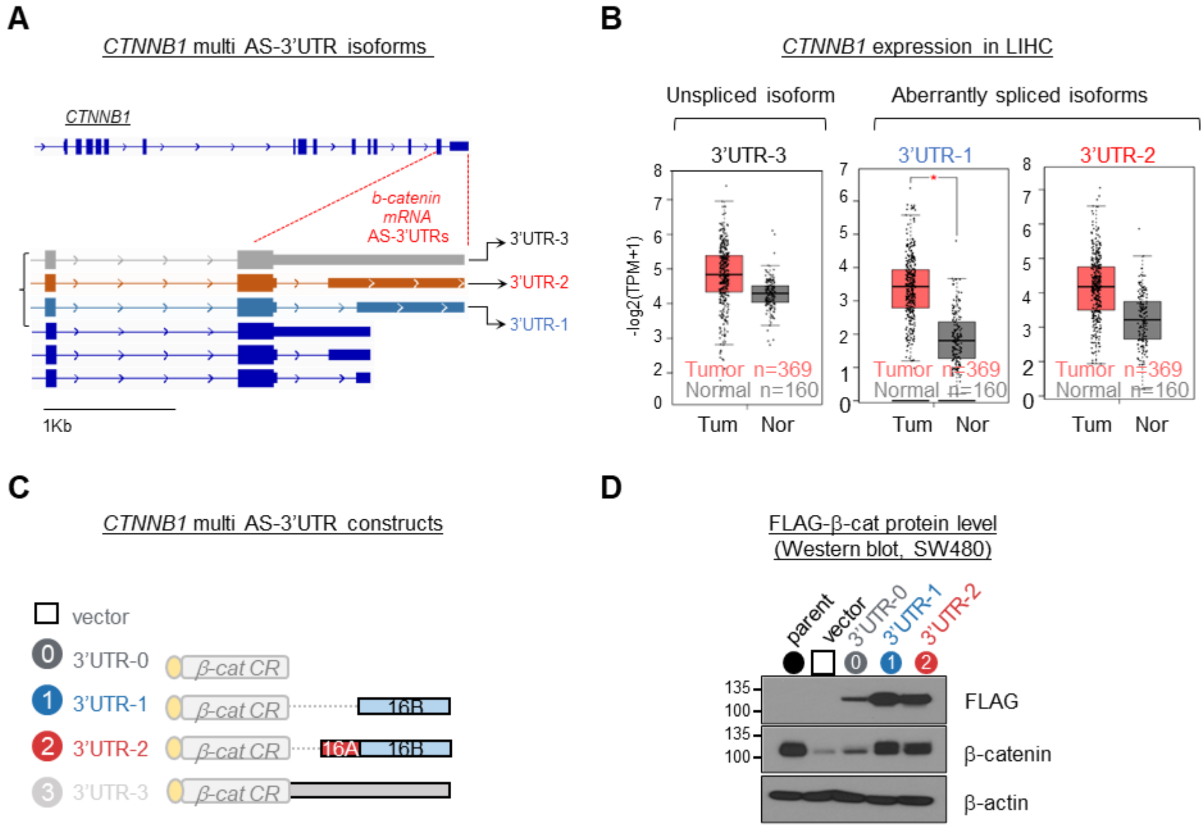


**Supplementary Figure 2. Establishment of SW480 and HT-29 stable cell lines expressing FLAG-β-catenin with distinct 3'UTRs (A)** A schematic diagram of the multiple *CTNNB1* mRNA isoforms with distinct 3'UTRs. **(B)** Expression profiles of CTNNB1 3'UTR isoforms in Liver Hepatocellular Carcinoma (LIHC) patient samples and normal tissues analyzed using the GEPIA2 platform. **(C)** Construction of FLAG-β-catenin expression vectors harboring different 3'UTRs for stable cell line generation. **(D)** Western blot analysis of FLAG-β-catenin expression in SW480 stable cell lines.

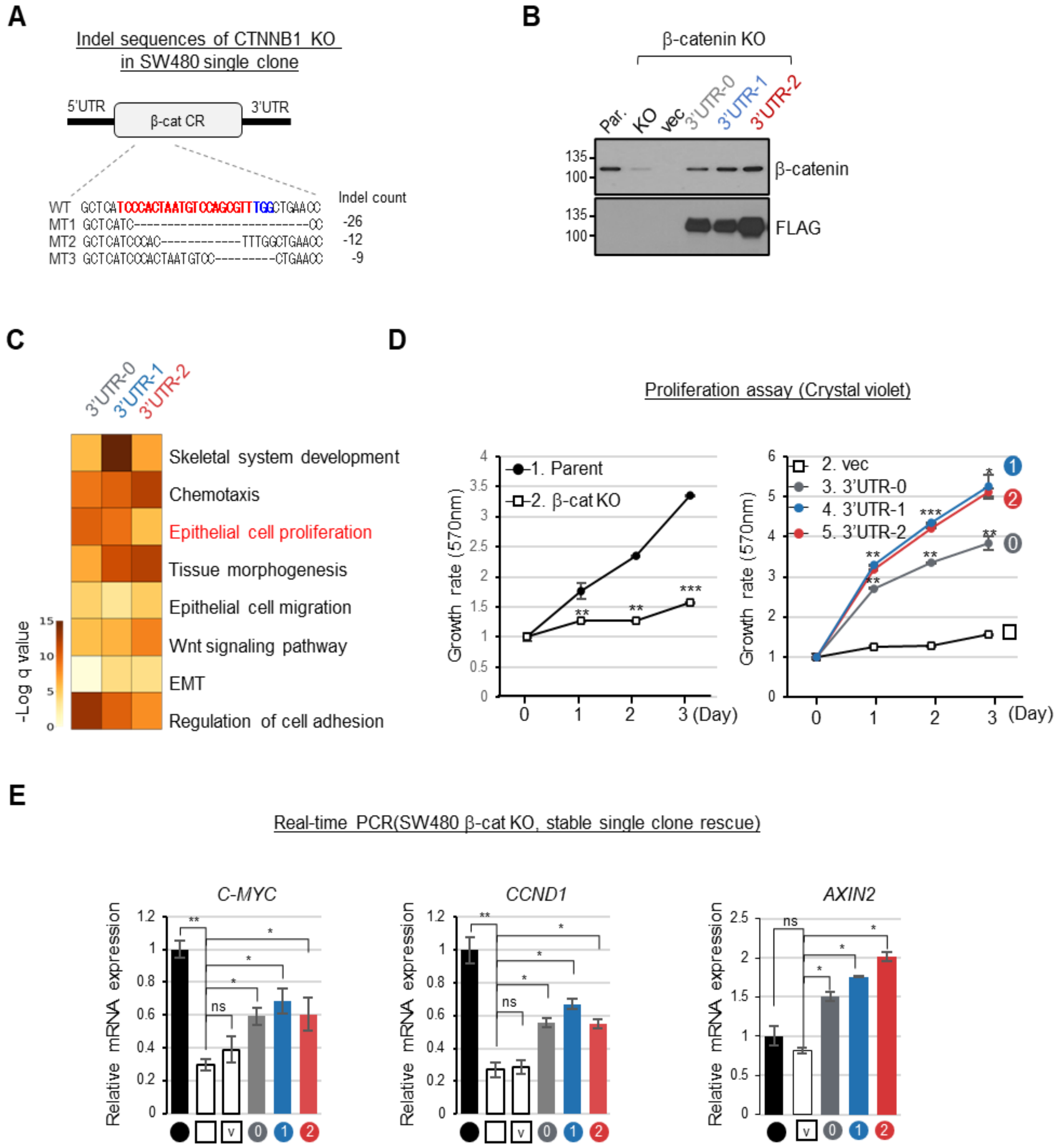


**Supplementary Figure 3. β-catenin-driven cell proliferation independent of 3'UTR isoforms (A)** DNA sequencing data showing CRISPR/Cas9-induced deletions in *CTNNB1* knockout clones. gRNA target sites highlighted in red. **(B)** Western blot confirming β-catenin knockout and FLAG-β-catenin expression from 3'UTR constructs in SW480. **(C)** Gene Ontology analysis of genes altered by β-catenin overexpression. **(D)** Cell proliferation assay by crystal violet staining of SW480 parental, knockout, and FLAG-β-catenin 3'UTRs rescue cells. **(E)** RT-qPCR analysis of canonical β-catenin target genes in FLAG-β-catenin 3'UTR-rescued SW480 cells.

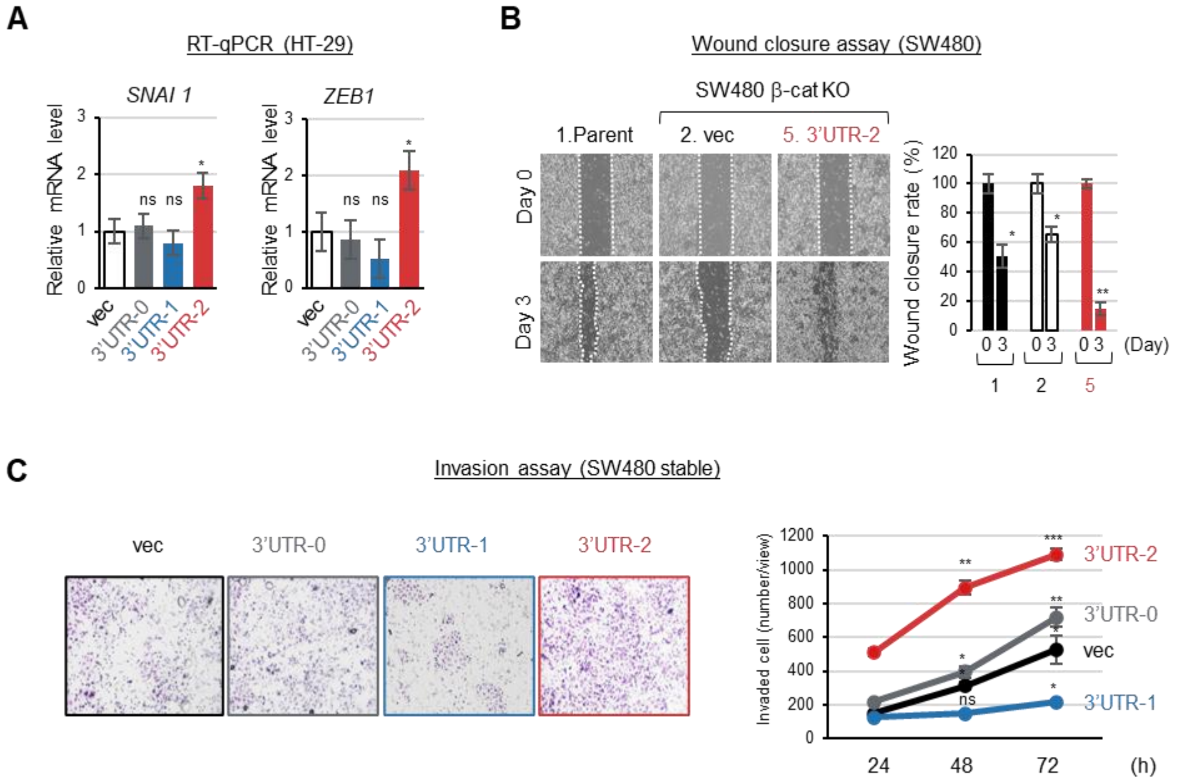


**Supplementary Figure 4. EMT-promoting β-catenin generated by aberrant 3'UTR**

**(A)** Validation of transcript level changes by RT-qPCR in HT-29 cells. **(B)** Wound closure assay of SW480 parental cells, SW480 cells expressing vec, and 3'UTR-2 derived β-catenin. **(C)** Transwell invasion assay of SW480 cells expressing vec, 3'UTR-0, 3'UTR-1, and 3'UTR-2-derived β-catenin cells seeded in Matrigel-coated inserts. Invaded cells at the lower surface were quantified after 24, 48, and 72 hours (n=3).

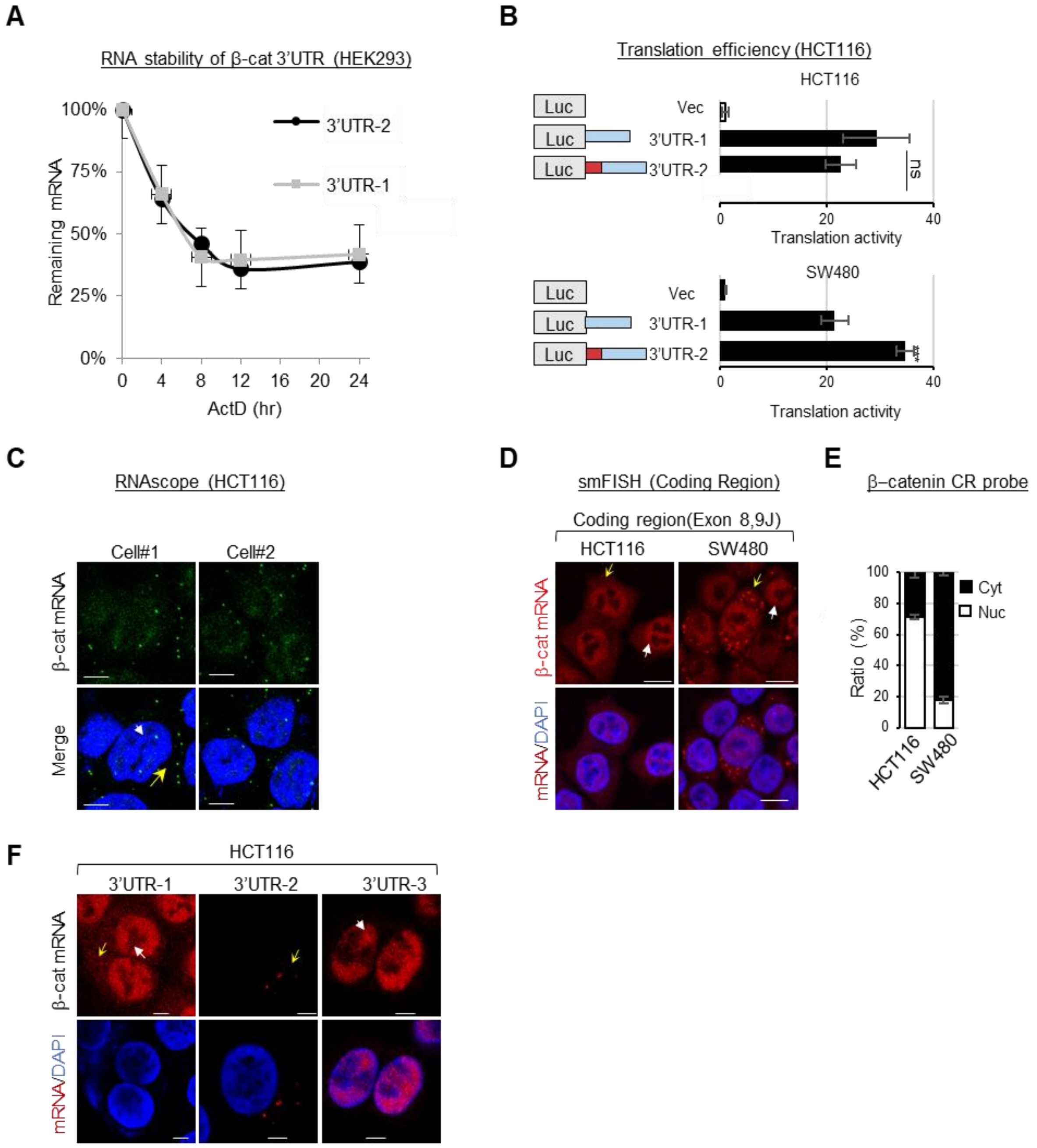


**Supplementary Figure 5. Functional characterization of β-catenin 3'UTR isoforms**

**(A)** mRNA stability assay showing the remaining levels of β-catenin mRNA containing 3'UTR-1 and 3'UTR-2 isoforms after actinomycin D treatment at the indicated time. **(B)** Luciferase reporter assay comparing β-catenin 3'UTR-1 and 3'UTR-2 isoforms in HCT116 (upper) and SW480 (lower) cells. **(C)** RNAscope images showing the distribution of total β-catenin mRNA in HCT116 cells. **(D, E)** smFISH analysis of β-catenin mRNA localization in HCT116 and SW480 cells **(D)**, and corresponding quantification of cytoplasmic and nuclear signal intensity **(E)**. **(F)** Representative smFISH images showing differential localization of β-catenin mRNA isoforms (3'UTR-1, 3'UTR-2, and 3'UTR-3) in HCT116 cells.

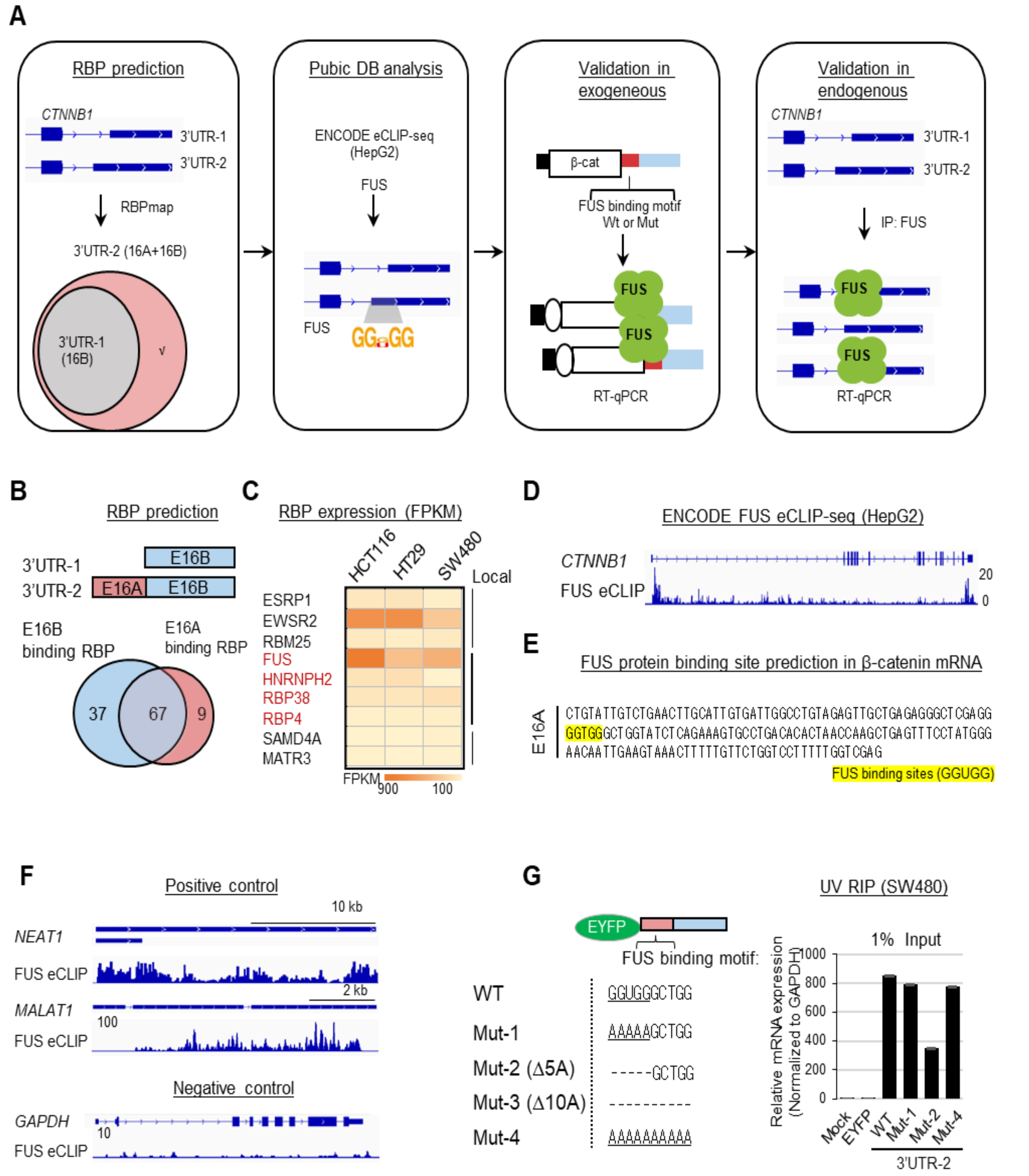


**Supplementary Figure 6. Identification of FUS binding motif in β-catenin 3'UTR**

**(A)** Workflow diagram outlining the strategy to identify RNA-binding proteins (RBPs) targeting alternative 3'UTR exons. **(B)** Venn diagram indicating the number of RBP candidates binding to alternative 3'UTR exons. **(C)** Heatmap of RBP candidates binding to exon 16A and their expression (FPKM) across CRC cell lines. **(D)** eCLIP-seq tracks showing FUS binding at the CTNNB1 locus in HepG2 cells. **(E)** Sequence of the β-catenin 3'UTR region from exon 16A to 16B with predicted FUS-binding site highlighted in yellow. **(F)** eCLIP-seq tracks showing FUS binding to established targets (NEAT1 and MALAT1), with no detectable binding at GAPDH, serving as positive and negative control. **(G)** UV-RNA-IP followed by RT-qPCR showing FUS association with chimeric 3'UTR reporters in SW480 cells. Expression of reporter constructs measured by RT-qPCR.

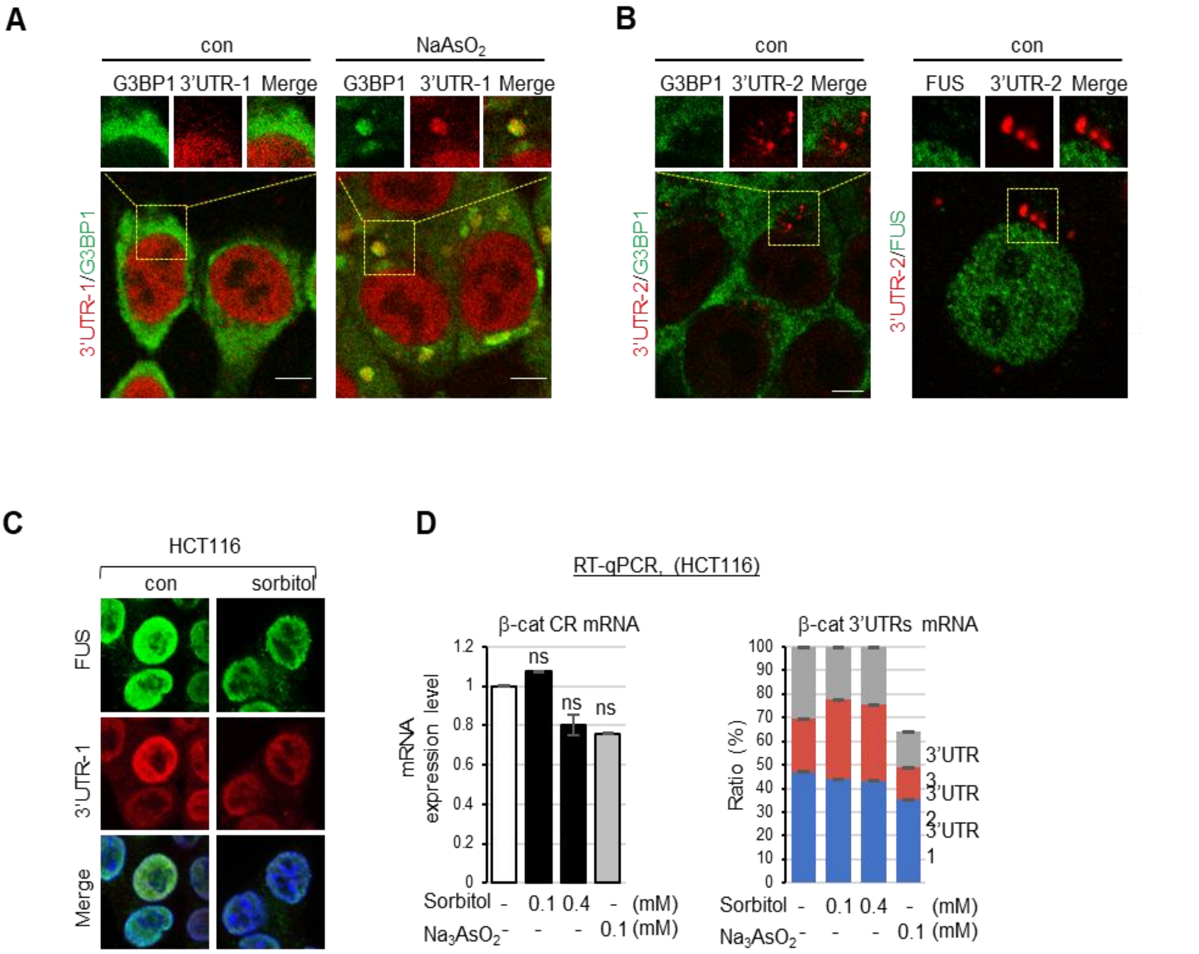


**Supplementary Figure 7. Stress-induced interaction between FUS and aberrant 3'UTR isoform**

**(A, C)** Dual smFISH and immunofluorescence analysis showing colocalization of β-catenin 3′UTR-1 mRNA with G3BP1 upon sodium arsenite treatment ($Na_3AsO_3$, 0.1 mM, 30 min) **(A)**, β-catenin 3′UTR-2 mRNA with G3BP1 (left) and FUS (right) **(B)**, and β-catenin 3′UTR-2 mRNA with FUS following sorbitol treatment (0.4 M, 30 min) **(C)**. **(D)** RT-qPCR analysis of β-catenin mRNA isoforms in HCT116 cells after sorbitol or $Na_3AsO_3$ treatments.

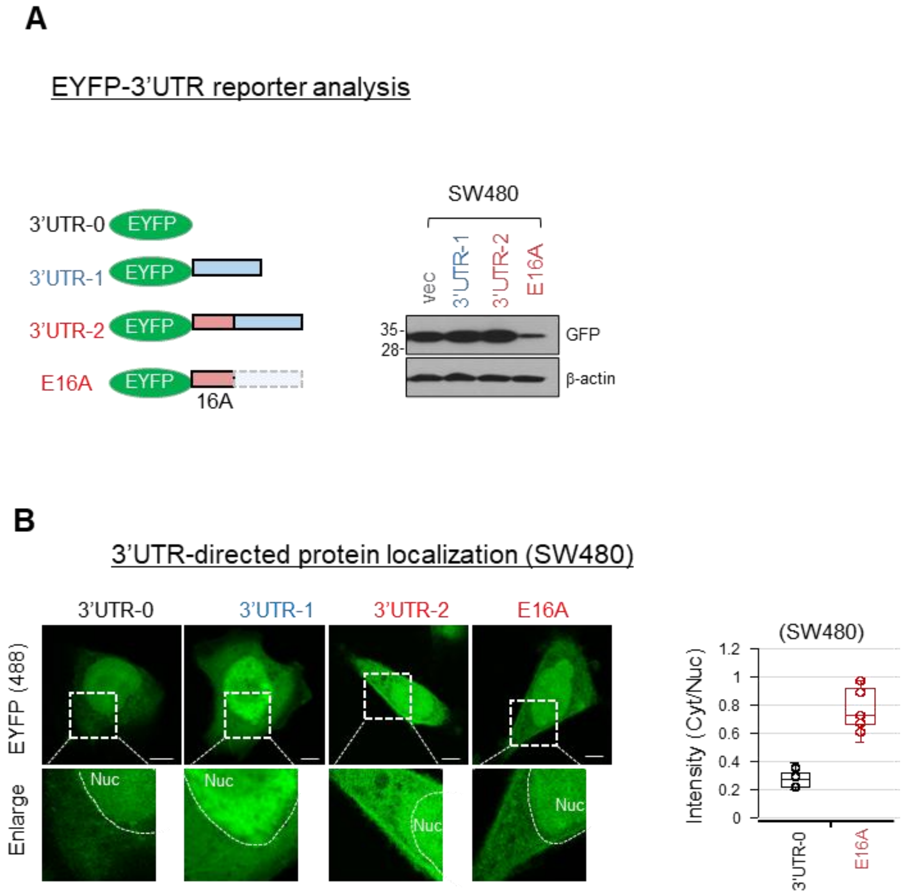


**Supplementary Figure 8. Functional analysis of β-catenin alternative 3'UTRs**

**(A)** Schematic and expression validation of EYFP constructs fused with alternative 3'UTR exons. **(B)** Immunofluorescence imaging of EYFP-3'UTR constructs in SW480 cells. Box plots showing the ratio of cytoplasmic to nuclear EYFP intensity.

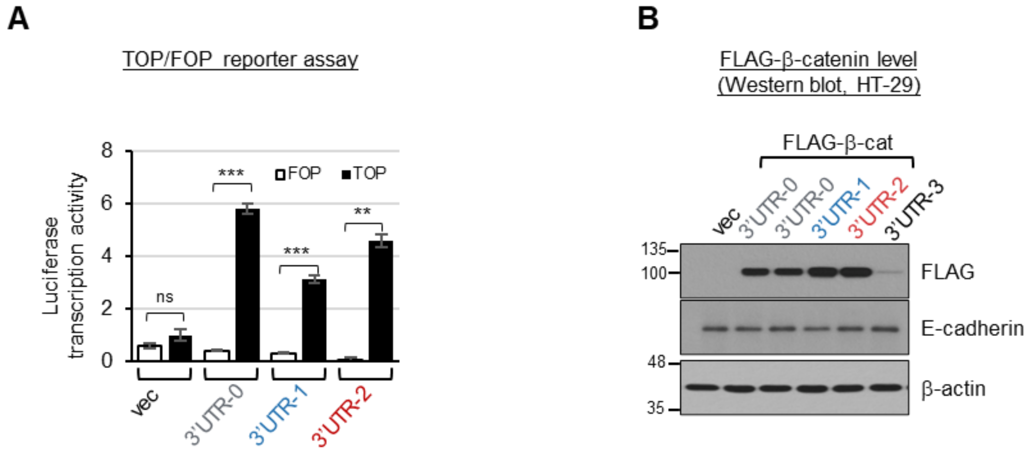


**Supplementary Figure 9. Functional validation of β-catenin 3′UTR-dependent transcriptional activity in stable cell lines. (A)** TOP/FOP luciferase reporter assays in the SW480 expressing β-catenin 3’UTR variants. **(B)** Western blot validation of FLAG-β-catenin expression in HT-29 stable cell lines.
